# Vision–language representations link food perception to subjective valuation

**DOI:** 10.64898/2026.11.01.763512

**Authors:** Yuhi Aoki, Asako Toyama, Michiyo Sugawara, Hiroki Kojima, Yuichi Yamashita, Shinsuke Suzuki

## Abstract

Subjective values of foods guide dietary decisions. Yet, how the brain computes such values from visual input remains unclear. Here, we show that visual food valuation relies on higher-level conceptual representations captured by a multimodal vision-language deep neural network model. Using food images and human ratings (N = 194 and 200), we found that a vision-language model predicted subjective value more accurately than vision-only models. Representational analyses demonstrated that, compared with the vision-only model, the vision-language model preferentially encoded higher-level food attributes, including perceived healthiness, while encoding less image-level visual information. Multivariate analyses of fMRI data (N = 31) showed that embedding features from both models corresponded to activity patterns along the visual pathways. Critically, vision-language features showed greater correspondence than vision-only features in the fusiform gyrus, a downstream region of the ventral visual pathway. These findings suggest that higher-level conceptual representations link visual food processing to subjective valuation.

## Introduction

Dietary decisions rely on the subjective values people assign to food items^1,2^. These values integrate multiple attributes, including anticipated taste, perceived healthiness, nutritional quality and visual appearance^3–8^. Food valuation has important implications for health, as maladaptive valuation processes have been implicated in obesity and eating disorders^9,10^. In everyday life, people often construct food values from visual cues before consumption. For example, when shopping for groceries, they visually inspect food products on supermarket shelves and infer likely taste, healthfulness and other desirable properties. Yet, little is known about how the brain transforms visual food information into subjective value^11^.

Accumulating evidence in decision neuroscience over the past two decades has identified a network of brain regions that signals the subjective value of food rewards, including the medial prefrontal cortex and ventral striatum^4,12–14^. These findings identify *where* the brain represents food value. They do not, however, explain *how* visual properties of food stimuli give rise to the representations that support valuation^11^. Thus, while existing research has characterized neural correlates of food value, it has not specified the representational processes that link visual food cues to subjective valuation.

Visual food valuation poses a representational problem: how does the appearance of a food item give rise to expectations about properties such as taste and healthiness that can inform its subjective value? Identifying these attributes as determinants of value does not, by itself, explain how they are inferred from an image. Addressing this question requires computational models that derive candidate representations from visual input, allowing their content and relevance to human valuation to be tested^11^.

Deep neural networks provide a computational framework for this approach by transforming images through successive layers of learned representations^15,16^. These intermediate representations can be examined for the information they encode and compared with both behavioral judgments and neural activity^17–20^. Comparing networks trained with different objectives further allows us to test which learned representations best account for human behavioral and neural responses^21–23^. For instance, this approach has been used to investigate visual and semantic contributions to human perception and memory^24,25^. That is, it may offer a way to formulate and evaluate representational accounts of visual food valuation.

Here, we compared representations in deep neural networks learned through image classification with those learned through image–language alignment. Image classification provides a baseline for the food information captured by representations optimized for visual recognition (e.g., convolutional neural networks, CNN)^26–28^. Vision–language training further aligns image representations with natural-language descriptions (e.g., contrastive language–image pretraining, CLIP)^29,30^. For example, an image of an apple and the text “apple” are mapped onto nearby locations in a shared representational space. This alignment may yield conceptual representations that capture both visual features and semantic information relevant to food valuation. We therefore asked whether a vision–language model more strongly represents food attributes such as taste, processing state and perceived healthiness, and whether its representations better account for human subjective value judgments.

Our previous work demonstrated that multimodal vision–language embeddings in CLIP can predict individual food evaluations^31^. However, predictive accuracy alone does not identify which properties of these representations are relevant to valuation. It also left open whether and how their organization corresponds to representations in the human brain. A representational account therefore requires in-depth examinations of what information the models encode, how that information varies across processing layers, and where the resulting representations correspond to neural activity during valuation.

We addressed these questions by combining deep neural network modeling, behavioral ratings and functional MRI. We first compared the ability of vision-language and vision-only representations to predict human subjective food value. We then employed representational similarity analysis to characterize the information encoded by each model and examined how this information varied across layers of the vision-language model. Finally, we tested how model representations corresponded to neural activity during food valuation, including whether vision-language representations explained neural representational structure beyond that captured by the vision-only model. We showed that vision-language representations predicted subjective food value more accurately in both lean and obese individuals. The representations encoded nutritive attributes, taste and perceived healthiness more strongly than vision-only representations. Layer-wise analyses further linked shallower representations to visual attributes and early visual cortex, and deeper representations to higher-level food attributes and downstream cortical regions, suggesting that higher-level food representations may link visual input to subjective valuation.

## Results

### Prediction of subjective food value by vision-language and vision-only models

We first tested whether vision-language representations predict subjective value ratings for food images more accurately than vision-only representations (Fig. 1a). We analyzed data from two online studies conducted by our laboratory: Online 1^31^ and Online 2^32^. In the datasets, lean and obese participants rated the subjective values of 896 food images (see Methods for details and Fig. S1a for histograms of the ratings; Online 1: *N* = 106 lean and 88 obese; Online 2: *N* = 107 lean and 93 obese).

**Figure 1:**
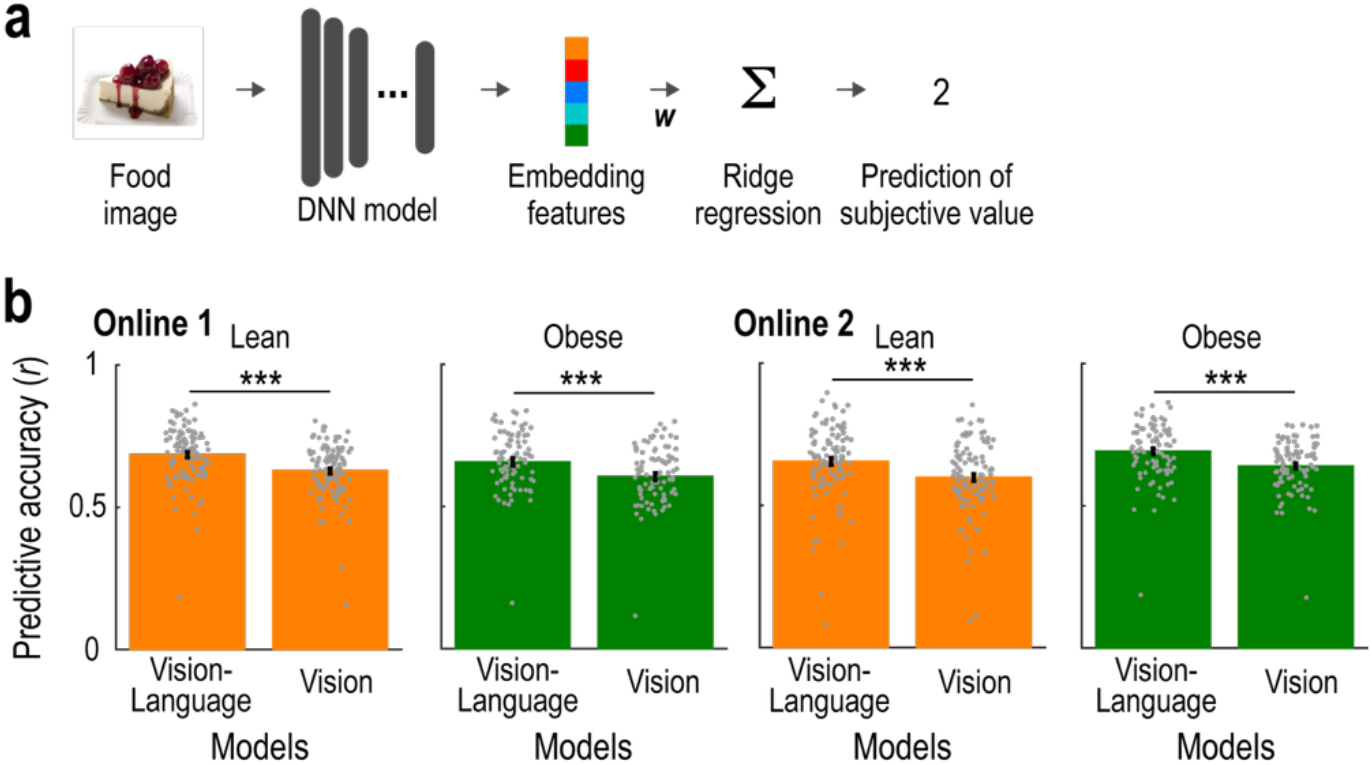
Prediction performance for subjective food value. (a) Schematic illustration of the prediction analysis. For each of the vision-language and vision-only deep neural network (DNN) models, we used ridge regression to predict human subjective value ratings from embedding features extracted from the output layer of the model. ***w***, weights of the embedding features. (b) Prediction performance. Predictive accuracy was quantified as the cross-validated correlation between predicted and observed ratings (mean ± SEM across participants). Each grey dot represents an individual participant. \*\*\**p* < 0.001, *t*-test (two-tailed).

As a vision-only model, we used ConvNeXt, a convolutional neural network trained on image classification^26^. As a vision-language model, we used CLIP, a multimodal deep neural network trained to align images and natural language in a shared representational space^30^. Notably, the vision-language model used ConvNeXt as its image encoder, enabling a direct comparison between vision-only and vision-language models with the visual backbone held constant. For each model, we extracted embedding features (i.e., activity pattern of the output layer) for each food image. These features were then fed into a ridge regression to predict subjective value rating. We evaluated prediction accuracy as the correlation between predicted and observed ratings using 5-fold cross-validation. Both models predicted subjective value ratings above chance across the two datasets and participant groups (Vision-language model: in Online 1, *t*(105) = 50.191 and *p* < 0.001 for the lean participants, and *t*(87) = 38.462 and *p* < 0.001 for the obese participants; in Online 2, *t*(106) = 37.554 and *p* < 0.001 for the lean participants, and *t*(92) = 45.968 and *p* < 0.001 for the obese participants; Vision-only model: in Online 1, *t*(105) = 50.382 and *p* < 0.001 for the lean participants, and *t*(87) = 37.937 and *p* < 0.001 for the obese participants; in Online 2, *t*(106) = 37.196 and *p* < 0.001 for the lean participants, and *t*(92) = 50.555 and *p* < 0.001 for the obese participants).

Crucially, the vision-language model predicted subjective value ratings more accurately than the vision-only model in both lean and obese participants across the two datasets (Fig. 1b). The prediction accuracies were significantly greater for the vision-language model than for vision-only model (Online 1: lean, *t*(105) = 21.216, *p* < 0.001; obese, *t*(87) = 18.595, *p* < 0.001; Online 2: lean, *t*(106) = 18.454, *p* < 0.001; obese, *t*(92) = 17.640, *p* < 0.001). Furthermore, we confirmed that these results held when we extracted vision-language embedding features from the same position within the image encoder as the vision-only features (see Methods and SI Text 1 for details). This advantage did not depend on the choice of vision-only model: the vision-language model also outperformed two additional vision-only models, ResNet^27^ and VGG16^28^, in both participant groups and datasets (Fig. S1b; comparison with ResNet in Online 1: lean, *t*(105) = 22.030, *p* < 0.001; obese, *t*(87) = 19.918, *p* < 0.001; Online 2: lean, *t*(106) = 18.235, *p* < 0.001; obese, *t*(92) = 15.686, *p* < 0.001; comparison with VGG16 in Online 1: lean, *t*(105) = 36.310, *p* < 0.001; obese, *t*(87) = 28.859, *p* < 0.001; Online 2: lean, *t*(106) = 27.173, *p* < 0.001; obese, *t*(92) = 28.209, *p* < 0.001). Together, these results demonstrated that vision-language representations capture information relevant to subjective food value beyond that captured by vision-only representations.

### Representational contents in the vision-language and vision-only models

Why did the vision-language model provide better prediction of subjective food value than the vision-only model? To answer the question, we sought to characterize the representational content of the vision-language model with comparison to the vision-only model, by using representational similarity analysis (RSA; see Fig. 2a and Methods)^33^. For each model, we constructed a representational dissimilarity matrix (RDM) from the embedding features (i.e., activity pattern of the output layer), quantifying how dissimilar each pair of food images was in the model representational space. We also constructed attribute-based RDMs from visual, nutritional and conceptual attributes of the food images. We then computed Spearman’s rank correlations between the model-based and attribute-based RDMs to assess the extent to which each model encoded information about each food attribute.

**Figure 2:**
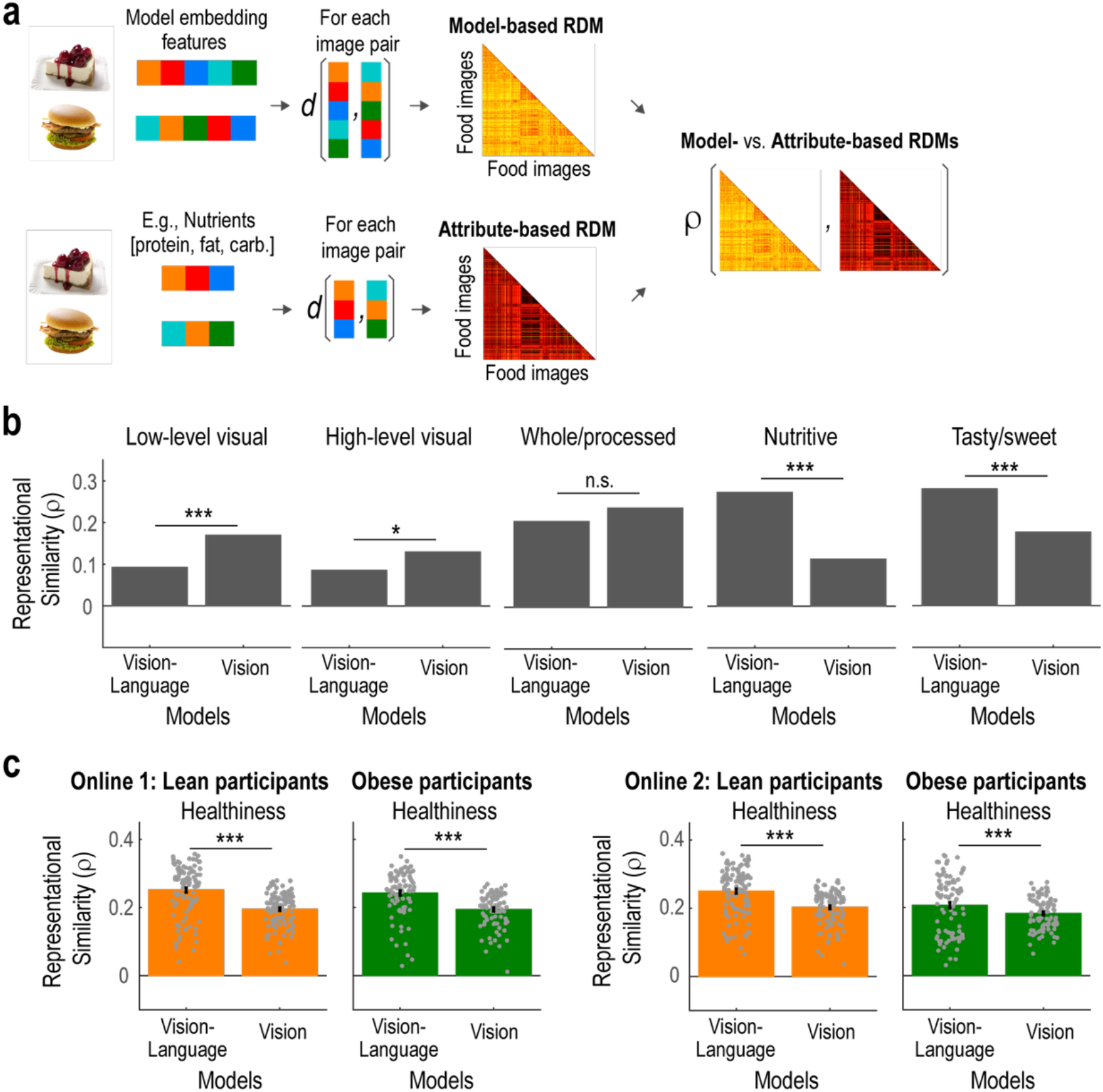
Representational content of the vision-language and vision-only models. (a) Schematic illustration of the representational similarity analysis (RSA). We constructed RDMs that represent the dissimilarity between pairs of food images in terms of model embedding features (model-based RDM) and food attributes such as nutrients (attribute-based RDM). We then compared these RDMs to assess the representational content of the models. *d*, distance between two vectors. RDM, representational dissimilarity matrix. (b) Representation of visual, nutritional, taste and processing-state attributes. We plotted representational similarity, quantified as Spearman’s rank correlation (*ρ*), between the model embedding features and each food attribute. n.s., not significant; \**p* < 0.05 and \*\*\**p* < 0.001, bootstrap test. (c) Representation of perceived healthiness. We plotted representational similarity between the model embedding features and perceived healthiness ratings (mean ± SEM across participants). Each grey dot represents an individual participant. \*\*\**p* < 0.001, *t*-test (two-tailed).

The vision-language model encoded visual attributes of the food images to a lesser extent than the vision-only model (Fig. 2b). Specifically, Spearman’s rank correlations were significantly weaker for the vision-language model for both low-level visual attributes (red, green and blue intensity values) and high-level visual attributes (local contrast, complexity and spatial frequency) (see Methods for details; low-level visual: *p* < 0.001, bootstrap test; high-level visual: *p* = 0.012, bootstrap test). Thus, compared with the vision-only model, the vision-language model less encoded visual attributes of the food images.

In contrast, the vision-language model encoded higher-level conceptual food attributes more strongly than the vision-only model. Spearman’s rank correlations were significantly greater for the vision-language model than for the vision-only model for nutritional attributes (protein, fat and carbohydrate contents) and taste (tasty or sweet) (Fig. 2b; nutritional attributes: *p* < 0.001, bootstrap test; and taste: *p* < 0.001, bootstrap test). We also examined perceived healthiness by correlating the model-based RDMs with RDMs derived from human ratings of healthiness. This analysis showed that the vision-language model encoded perceived healthiness more strongly than the vision-only model across both datasets and participant groups (Fig. 2c; Online 1: lean, *t*(105) = 11.418, *p* < 0.001; obese, *t*(87) = 10.453, *p* < 0.001; Online 2: lean, *t*(106) = 9.542, *p* < 0.001; obese, *t*(92) = 3.997, *p* < 0.001). Collectively, these representational analyses show that, compared with the vision-only model, the vision-language model preferentially encoded higher-level food attributes, including nutritional attributes, taste and perceived healthiness, while encoding less image-level visual information.

### Neural representations of the embedding features in the vision-language model

We have so far shown that embedding features (i.e., activity pattern of the output layer) from the vision-language model predicted subjective food value more accurately than those from the vision-only model, while also encoding higher-level food attributes more strongly. We next asked whether and how these embedding features corresponded to neural activity patterns elicited by food images.

To this end, we scanned 31 participants with functional MRI while they rated the subjective values of 568 food images over three days (Fig. 3a)^34^. The behavioral data replicated the findings in the online datasets. That is, embedding features from the vision-language model predicted subjective value ratings more accurately than those from the vision-only model (Fig. S1b; t(30) = 11.236, p < 0.001). Furthermore, the vision-language model outperformed other vision-only models (Fig. S1b; comparison with ResNet: t(30) = 8.323, p < 0.001; and comparison with VGG16: t(30) = 14.175, p < 0.001).

**Figure 3:**
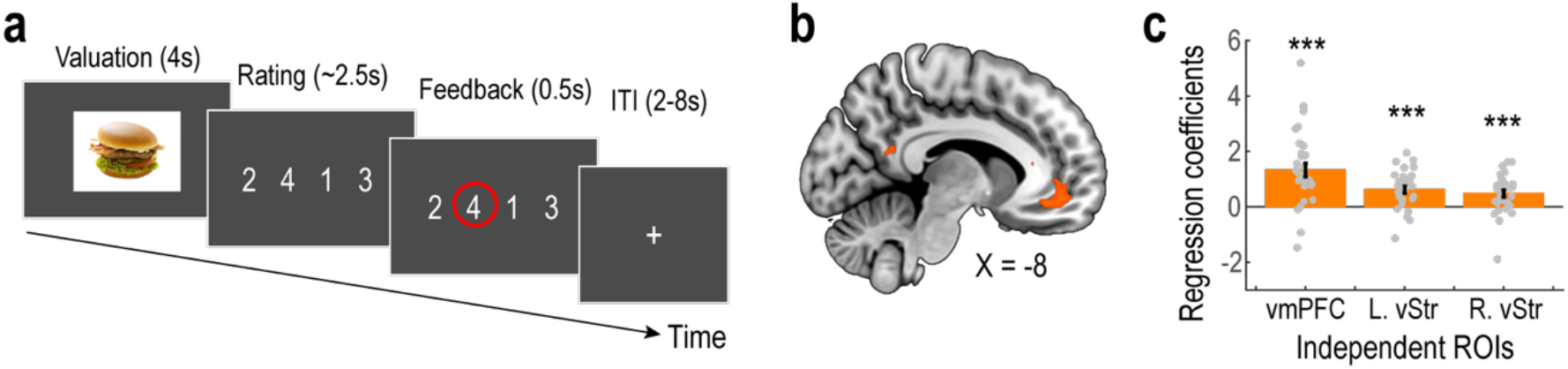
fMRI experiment and basic results. (a) Trial timeline. On each trial, participants rated the subjective value of a food item. In the rating phase, the mapping between response keys and rating values was randomized across trials. (b) Whole-brain univariate analysis of the subjective value signal. The statistical parametric map shows fMRI signals parametrically modulated by subjective value ratings during the valuation phase. The map is thresholded at *p* < 0.05, voxel-level FWE corrected. (c) Independent region-of-interest (ROI) analysis of the subjective value signal. Bar plots show regression coefficients for subjective value (mean ± SEM across participants). Each grey dot represents an individual participant. \*\*\**p* < 0.001. vmPFC, ventromedial prefrontal cortex. L. vStr, left ventral striatum. R. vStr, right ventral striatum.

A whole-brain univariate analysis of the fMRI data confirmed a well-established association between subjective value and activity in the ventromedial prefrontal cortex (vmPFC) at the time of valuation. Specifically, BOLD signals in the vmPFC during the valuation phase were significantly correlated with the subjective value of the food item presented on each trial (Fig. 3b; *p* < 0.05, voxel-level FWE corrected). Independent region-of-interest analyses further showed that BOLD signals in reward-processing regions, including the vmPFC and bilateral ventral striatum, were positively associated with subjective value (Fig. 3c; vmPFC: *t*(30) = 5.472, *p* < 0.001; left ventral striatum: *t*(30) = 5.148, *p* < 0.001; right ventral striatum: *t*(30) = 3.799, *p* < 0.001).

We next employed multivoxel pattern analyses to identify brain regions where embedding features (activity pattern of the output layer) from the vision-language and vision-only models corresponded to neural activity patterns elicited by food images (Fig. 4a). We constructed fMRI-based RDMs that quantified the dissimilarity between each pair of food images in multivoxel neural activity patterns^33^. We then correlated these fMRI-based RDMs with model-based RDMs derived from the embedding features of each model. This RSA identified brain regions where neural activity patterns corresponded to embedding features from the vision-language and vision-only models, respectively.

**Figure 4:**
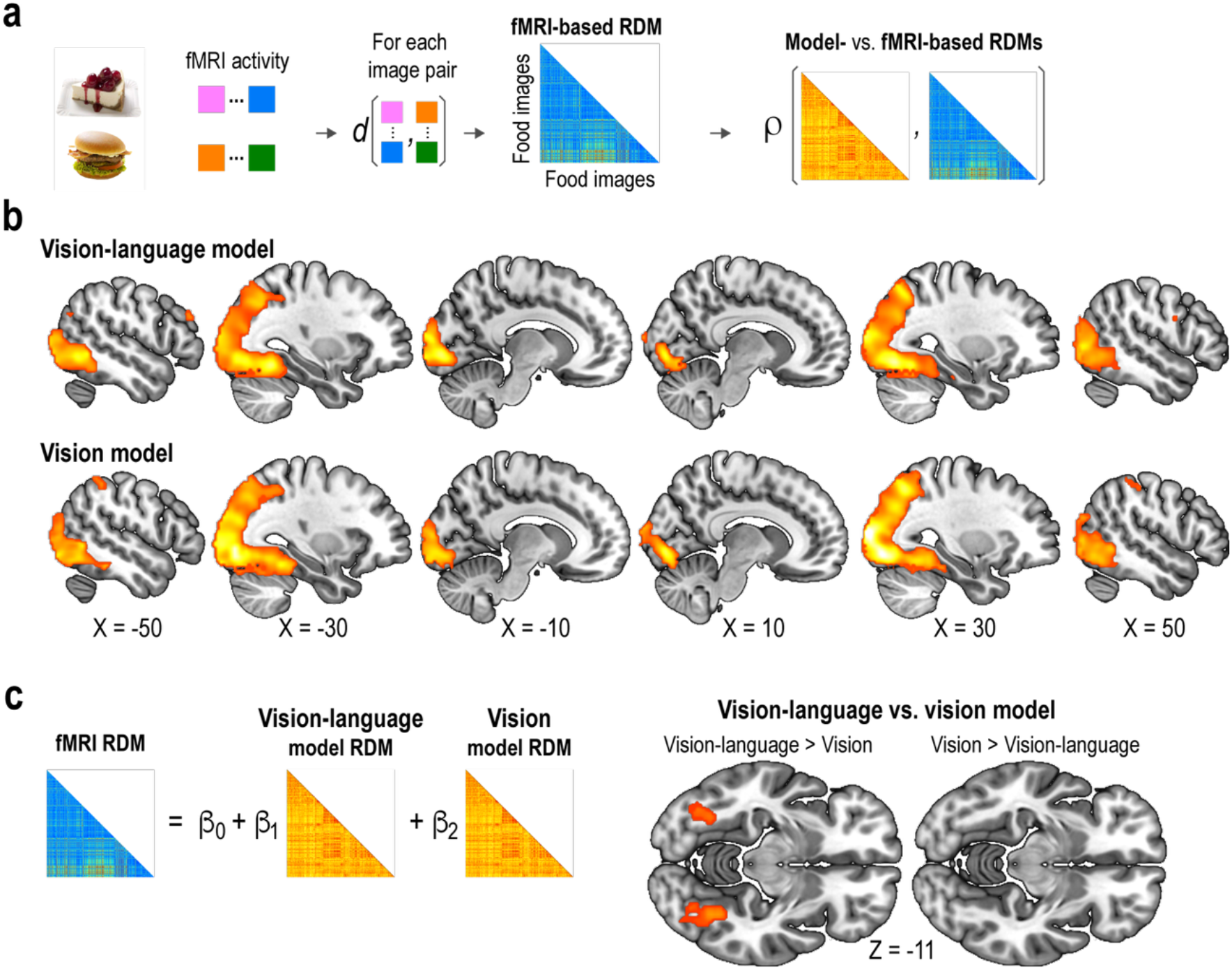
Neural representations of model embedding features. (a) Schematic illustration of the representational similarity analysis (RSA). We constructed RDMs that represent the dissimilarity between pairs of food images in terms of fMRI responses (fMRI-based RDM). We then compared the RDMs with model-based RDMs (see Fig. 2a) to assess the representational correspondence between the models and neural activity. RDM, representational dissimilarity matrix. (b) Whole-brain searchlight RSA of the vision-language and vision-only models. Statistical parametric maps show representational similarity between model embedding features and multivoxel fMRI activity patterns during the valuation phase. The maps are thresholded at *p* < 0.05, voxel-level FWE corrected. (c) Whole-brain searchlight RSA of the contrast between vision-language and vision-only models. Schematic illustration depicts the multiple-regression RSA. Statistical parametric maps show the contrasts in the representational similarity between vision-language and vision-only model embedding features and fMRI activity patterns during the valuation phase. The maps are thresholded at *p* < 0.05, voxel-level FWE corrected.

Whole-brain searchlight RSA revealed that, during valuation, the representational geometry of embedding features from the vision–language model corresponded to multivoxel fMRI activity patterns throughout the visual pathways (Fig. 4b and Table S1; voxel-level FWE-corrected *p* < 0.05). Significant effects extended from the primary visual cortex into downstream regions of the dorsal and ventral visual pathways, including the inferior parietal lobule and fusiform gyrus. Embedding features from the vision-only model also corresponded to multivoxel activity patterns along the visual pathways (Fig. 4b and Table S1; voxel-level FWE-corrected *p* < 0.05). To directly compare the neural correspondence of the two models, we next conducted multiple-regression RSA^35,36^, in which we regressed fMRI-based RDMs on the two RDMs derived from the vision-language and vision-only models respectively. Whole-brain searchlight analysis revealed that vision-language embedding features showed significantly greater correspondence with fMRI activity patterns than vision-only embedding features in the bilateral fusiform gyrus (Fig. 4c and Table S1; voxel-level FWE-corrected p < 0.05).

Anatomically defined ROI RSA yielded results consistent with the whole-brain searchlight analysis (Fig. S2). Specifically, in a direct within-participant comparison, correspondence with multivoxel fMRI activity patterns was significantly greater for the vision–language model than for the vision-only model in ventral visual pathway regions (VO, PHC and LO) and downstream ventral regions, including the bilateral fusiform gyri (Fig. S2c; *p* < 0.05, Bonferroni-corrected across 26 bilateral ROIs, two-tailed t-test). Together, these findings indicate that vision–language embedding features captured neural representational structure along the ventral visual pathway, extending into higher-order downstream regions beyond the correspondence captured by vision-only embedding features.

### Layer-wise representational contents in the vision-language model and the neural correspondences

We have so far examined representational contents in the embedding features (i.e., activity pattern of the output layer). Next, we asked how visual and higher-level food attributes were represented across layers of the vision-language model. Applying RSA separately to each layer of the vision-language model, we found that visual attributes corresponded most strongly to representations in the first and second layers (Fig. 5a). By contrast, deeper layers, but not shallow layers, represented higher-level food attributes, including nutritional attributes, taste and processing state (Fig. 5a). Perceived healthiness and subjective value ratings from both lean and obese participants also corresponded more strongly to deeper layers (Fig. 5b). The representational pattern was preserved when model activations were extracted from each layer after normalization (Fig. S3; see Methods for details). These results suggest that the vision-language model progressively transforms low-level visual information into higher-level conceptual representations of food attributes.

**Figure 5:**
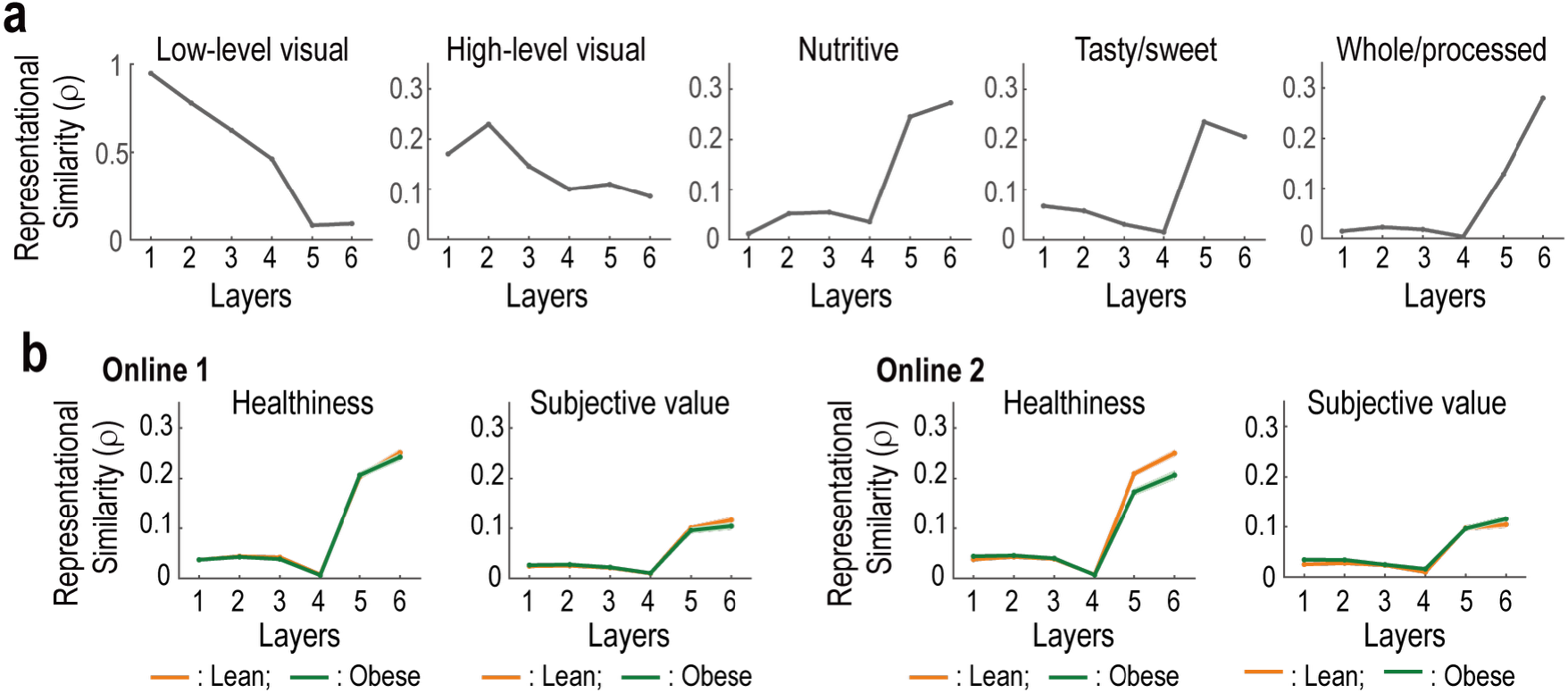
Layer-wise representations in the vision-language model. (a) Representation of visual, nutritional, taste and processing-state attributes across layers. For each layer, we plotted representational similarity, quantified as Spearman’s rank correlation (ρ), between the model embedding features and each food attribute. (b) Representation of perceived healthiness across layers. For each layer, we plotted representational similarity between the model embedding features and perceived healthiness ratings as well as subjective value (mean ± SEM across participants). *Left*, Online 1. *Right*, Online 2.

Finally, we examined where multivoxel fMRI activity patterns corresponded to layer-specific representations in the vision-language model and to subjective value. Whole-brain searchlight RSA was conducted separately for each model layer, model-predicted subjective value, and participants’ actual subjective value rating. Shallower model layers corresponded primarily to activity patterns in the early visual cortex (Fig. 6a and Table S2; *p* < 0.05, voxel-level FWE corrected). By contrast, correspondence with deeper layers extended beyond the early visual cortex to higher-order regions along the ventral and dorsal visual pathways and to lateral prefrontal regions (Fig. 6a and Table S2; *p* < 0.05, voxel-level FWE corrected). Model-predicted subjective value corresponded to activity patterns in higher-order visual regions as well as in lateral and dorsal prefrontal regions (Fig. 6a and Table S3; p < 0.05, voxel-level FWE corrected). Participants’ actual subjective value corresponded to activity patterns in the medial and ventrolateral prefrontal cortex (Fig. 6a and Table S4; p < 0.05, voxel-level FWE corrected).

**Figure 6:**
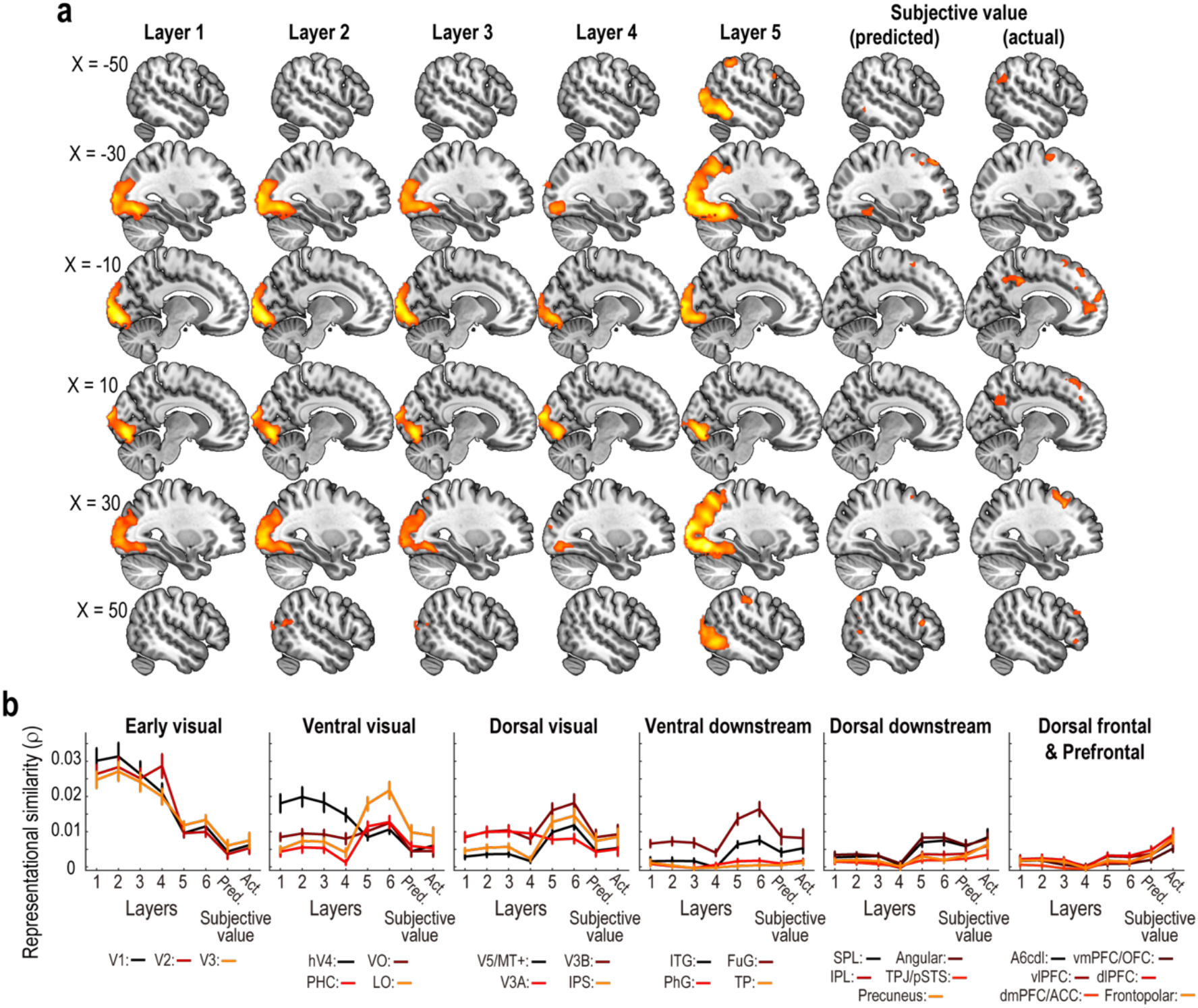
Layer-wise neural representations in the vision-language model. (a) Whole-brain searchlight representational similarity analysis (RSA) of each layer in the vision-language model and the subjective values. Statistical parametric maps show representational similarity between model embedding features and multivoxel BOLD activity patterns during the valuation phase. Layer 6 is not shown as the map is identical to Fig. 4b. The maps are thresholded at *p* < 0.05, voxel-level FWE corrected. (b) Anatomically defined ROI RSA of each layer in the vision-language model and the subjective values. Representational similarity between embedding features from each layer in the vision-language model and multivoxel fMRI activity patterns during the valuation phase is plotted for each anatomically defined region of interest (ROI). Points and error bars indicate the mean ± SEM across participants. V1, Primary visual cortex / primary visual area; V2, Secondary visual cortex / visual area 2; V3, Visual area 3; hV4, Human visual area 4; VO, Ventral occipital areas; PHC, Parahippocampal cortex; LO, Lateral occipital areas; ITG, Inferior temporal gyrus; FuG, Fusiform gyrus; PhG, Parahippocampal gyrus; Temporal pole, Temporal pole; V5/MT+, Visual area 5 / middle temporal visual complex; V3B, Visual area V3B; V3A, Visual area V3A; IPS, Intraparietal sulcus; SPL, Superior parietal lobule; Angular, Angular gyrus; IPL, Inferior parietal lobule; TPJ/pSTS, Temporoparietal junction / posterior superior temporal sulcus; A6cdl, Caudodorsal lateral area 6 (dorsal premotor cortex); vmPFC/OFC, Ventromedial prefrontal cortex / orbitofrontal cortex; vlPFC, Ventrolateral prefrontal cortex; dlPFC, Dorsolateral prefrontal cortex; dmPFC/ACC, Dorsomedial prefrontal cortex / anterior cingulate cortex; Frontopolar, Frontopolar cortex. Pred, predicted subjective value rating. Act., actual subjective value rating.

Anatomically defined ROI analyses revealed a more fine-grained pattern (Fig. 6b; see Fig. S4 for detailed statistics). Shallower model layers corresponded primarily to activity patterns in the early visual cortex (V1, V2, and V3). Correspondence with deeper layers extended to higher-order regions along the ventral and dorsal visual pathways (VO, PHC, LO, ITG, FuG, V5/MT+, V3B, V3A, and IPS) and to parietal association regions (SPL, angular gyrus, and IPL). Subjective value predicted from the final-layer embedding by the ridge-regression model corresponded to a distributed network encompassing higher-order visual, parietal, and lateral prefrontal regions, including PHC, LO, V5/MT+, the angular gyrus, precuneus, dlPFC, and frontopolar cortex. Participants’ actual subjective value corresponded additionally to medial and ventrolateral prefrontal regions, including the vmPFC/OFC, dmPFC/ACC, and vlPFC. The pattern of neural representations remained unchanged when we extracted model activations from each layer after normalization (Fig. S5 and S6; see Methods for details).

We next examined whether each layer showed distinct neural correspondence after controlling for the effects of the other layers. To this end, we conducted multiple-regression RSA^35,36^, in which we regressed fMRI-based RDMs on RDMs derived from layer-specific embedding features and subjective value (Fig. S7). Because RDMs from adjacent layers were highly correlated (Fig. S7a), we focused on the first, fourth and sixth layers (Fig. S7ab). Whole-brain searchlight RSA showed that embedding features from the first and fourth layers corresponded to BOLD activity patterns primarily in the early visual cortices (Fig. S7cd; p < 0.05, voxel-level FWE corrected). By contrast, embedding features from the deeper sixth layer corresponded not only to activity patterns in the early visual cortices, but also to those in higher-order downstream visual cortices and lateral prefrontal cortices (Fig. S7e; p < 0.05, voxel-level FWE corrected). Subjective value corresponded to BOLD activity patterns in the medial prefrontal cortex and ventrolateral prefrontal cortex (Fig. S7f; p < 0.05, voxel-level FWE corrected).

Together, these findings suggest a hierarchical representational pathway through which visual food information is transformed into higher-level conceptual representations that may support constructing subjective value signals in prefrontal valuation regions.

## Discussion

This study provides insight into the neurocomputational mechanisms underlying subjective food valuation. By comparing representations derived from deep neural network models with human subjective value ratings and fMRI data, we sought to elucidate how the brain transforms visual food information into subjective value. Specifically, we compared vision-only deep neural network models pretrained primarily for image classification to a multimodal vision-language model trained to align visual representations with linguistic and semantic information. We then examined the representational content of these models and their neural correlates.

We demonstrated that the multimodal vision-language deep neural network model predicted subjective food value more accurately than the vision-only model, even though both models used the ConvNeXt-Base backbone architecture. This finding suggests that subjective food value cannot be fully explained by representations acquired through image classification alone. Instead, representations aligned with semantic information provide additional explanatory power for predicting how humans evaluate food images. This interpretation is consistent with recent work showing that deep neural networks trained on images, text or image–text pairs can disentangle visual, visual–semantic and semantic components of human mental representations, and that integrated visual–semantic representations make a unique contribution to perception and memory^24,25,37^. More broadly, multimodal vision-language models have been shown to capture human-relevant visual–semantic representations across several domains, including image aesthetics^38^, material perception^39^, face-based social perception^40^ and food valuation^31^. The present study extends these findings by showing that the advantage of the vision-language model was consistently observed in both lean and obese participants across multiple datasets. This consistency supports the replicability and generalizability of the effect and further suggests that visual food valuation may depend on representations that integrate visual information with higher-level semantic structure.

More importantly, we identified representational content that may account for the advantage of the vision-language model. Compared with the vision-only model, embedding features (i.e., activity in the output layer) of the vision-language model preferentially encoded higher-level food attributes, including nutritional properties, taste and perceived healthiness, while encoding less image-level visual information. Thus, its superior performance in predicting subjective food value may reflect conceptual and semantic information that is not captured to the same extent by the vision-only model. Notably, the presence of semantic structure in the vision-language model does not necessarily imply that the model understands semantic meaning in the same way as humans do. Rather, our findings suggest that contrastive learning with language induces statistical structure in image embeddings that corresponds to attributes such as taste, healthiness and nutritional state. Taken together, these findings provide a mechanistic account of how vision-language models can predict human food valuation more accurately than vision-only models.

At the neural level, embedding features from the vision-language model (i.e., activity in its output layer) corresponded to multivoxel fMRI activity patterns along both the ventral and dorsal visual pathways^41,42^, extending into their downstream regions. In the ventral pathway, correspondence extended from human V4, ventral occipital, parahippocampal and lateral occipital visual areas to the inferior temporal gyrus, fusiform gyrus, parahippocampal gyrus and temporal pole. This spatial progression may reflect the extraction of food-related meanings and attributes from increasingly complex visual representations, ranging from shape and object-category information to higher-level conceptual information^43,44^. In particular, the extension into the fusiform and inferior temporal gyri suggests a contribution of higher-order object representations^43,45,46^. In the dorsal pathway, correspondence extended from dorsal extrastriate visual areas, including areas V3A, V3B and V5/MT+, through the intraparietal sulcus to the superior parietal lobule, inferior parietal lobule and angular gyrus. This pattern suggests that neural representations relevant to food valuation extend beyond the ventral visual pathway. These dorsal regions may contribute information related to visual attention, spatial processing, action affordances and sensorimotor transformations that support choice^47–50^. Importantly, in the bilateral fusiform gyrus, vision–language embedding features showed greater correspondence with fMRI activity patterns than vision-only features. This finding suggests that the representational structure of food images in the fusiform cortex is better captured by language-aligned visual embeddings than by image-classification-trained visual features, possibly reflecting visual–semantic representations that integrate higher-order object information with conceptually organized food attributes.

Furthermore, we examined representations across individual layers of the vision-language model. Shallower layers corresponded more strongly to visual attributes, whereas deeper layers corresponded more strongly to higher-level food attributes, including taste, processing state and perceived healthiness, as well as subjective value. These findings support a hierarchical computational process in which low-level visual information is progressively transformed into higher-level conceptual representations of food^11^. Analysis of the fMRI data revealed a parallel neural hierarchy: shallower model layers corresponded primarily to early visual areas, whereas deeper layers showed correspondence extending into higher-order and downstream regions of the ventral and dorsal visual pathways^18,19^, as well as the lateral prefrontal cortex. This pattern suggests a correspondence between the representational hierarchy of the artificial model and the hierarchy of visual processing in the human brain. Consistent with previous studies^12^, subjective value was represented in prefrontal regions, including the medial prefrontal cortex, indicating that activity patterns in these regions tracked the subjective values assigned to food images. Notably, deeper layers of the vision-language model corresponded to the lateral prefrontal cortex, whereas subjective value was represented in the medial and ventrolateral prefrontal cortices. Together, these findings raise the possibility that higher-level conceptual representations of food provide inputs to prefrontal computations of subjective value.

This study has several limitations. First, the result that the vision-language model predicted subjective value ratings more accurately than the vision-only model does not, by itself, establish that it relied on linguistic or semantic information. This possibility should be evaluated cautiously by integrating evidence from multiple complementary approaches. Second, the vision-language model evaluated in this study is based on a specific architecture, training dataset, and learning objective. Therefore, the differences with the vision-only model cannot be attributed solely to language-related information, as they may also reflect differences in factors such as contrastive learning, the scale of the training data, and the projection layers. Third, the generalizability of our findings to everyday decision-making involving actual food consumption, hunger states, choice, purchasing behavior, and contextual or social information is limited. In particular, although cross-cultural differences in food preferences are beyond the scope of the present study, they represent an important consideration. It should also be noted that the food images used in this study predominantly depicted Western foods^51^. Given the substantial regional and cultural diversity in food preferences, future studies should examine these findings across diverse populations and regions worldwide to assess their generalizability.

In conclusion, by combining multimodal deep neural network modelling with functional MRI, we propose a neurocomputational framework in which language-aligned higher-order representations of food attributes bridge visual input and subjective value. Beyond localizing subjective value signals to specific brain regions, the present study sheds light on how such value representations are constructed from visual food information in the brain. Given the close relationship between subjective food preferences and eating disorders, our findings may also have implications for understanding the pathophysiology of eating disorders and, ultimately, for developing more effective interventions.

## Methods

We analyzed the *three* datasets collected in our laboratory: Online 1^31^, Online 2^32^, and fMRI^34^. This study was approved by the Ethics Committee of National Center of Neurology and Psychiatry (A2021-072 for Online 1 experiments) and by the Hitotsubashi University IRB Committee (2024C030 for Online 2 and 2025C069 for fMRI experiments). All participants provided their informed consent.

### Participants

Online 1 included 194 participants. The lean group consisted of 106 participants (57 females and 49 males; age = 38.31 ± 10.04 years [mean ± SD]; BMI = 20.91 ± 2.11 kg/m²), and the obese group consisted of 88 participants (49 females and 39 males; age = 39.90 ± 12.70 years; BMI = 32.83 ± 3.25 kg/m²). Online 2 included 200 participants. The lean group consisted of 107 participants (53 females and 54 males; age = 39.75 ± 10.53 years; BMI = 20.57 ± 1.14 kg/m²), and the obese group consisted of 93 participants (45 females and 48 males; age = 40.16 ± 10.56 years; BMI = 32.59 ± 3.09 kg/m²). The fMRI experiment included 31 participants (11 females and 20 males; age = 21.29 ± 1.87 years; BMI = 20.63 ± 2.64 kg/m²). All participants were native Japanese speakers. In the fMRI experiment, participants were pre-screened to exclude individuals with a history of neurological or psychiatric illness. See the references^31,32,34^ for details.

### Stimuli

We used food images from the *food-pics extended* database, which contains 896 images along with information on their visual, nutritive and higher-order attributes^51^.

### Experimental procedure

#### Online 1 and 2

Participants completed an online food evaluation task implemented in Qualtrics (https://www.qualtrics.com/) over three separate days within a 10-day period. On each day, they evaluated all 896 food images on one of three dimensions: subjective value, tastiness or perceived healthiness. Ratings were provided on an eight-point scale ranging from 1 (strongly disagree) to 8 (strongly agree), and the order of the three tasks was randomized across participants. They were instructed to refrain from consuming food or beverages other than water for 3 hours before each session. Further procedural details are provided in the references^31,32^.

#### fMRI

Participants performed a food valuation task during fMRI scanning on three separate days (Fig. 3a). The task comprised 568 food images presented in randomized order across 12 runs, with four runs per day and 47 or 48 trials per run. Participants were instructed to refrain from consuming food or beverages other than water for 3 h before each scanning session. On each trial, a food image was presented for 4 s, during which participants evaluated its subjective value (valuation phase). They then reported their evaluation using a four-point scale, with higher scores indicating greater subjective value, by pressing the corresponding keypad button within 2.5 s (rating phase). The selected rating was subsequently displayed for 0.5 s (feedback phase).

Inter-trial intervals were sampled from a uniform distribution between 2 and 8 s. Participants were instructed to form their evaluation during the valuation phase. To dissociate subjective value from response location and motor preparation, the mapping between rating values and keypad positions was randomized across trials. Further details are provided in the reference^34^.

### Behavioral data analysis

#### Prediction of subjective food value using deep neural networks

We predicted subjective food value from image features extracted from pretrained vision–language and vision-only deep neural networks. The vision–language model was CLIP with a ConvNeXt-Base visual backbone, implemented in OpenCLIP using the pretrained checkpoint laion400m_s13b_b51k^29,30^. The primary vision-only model was ConvNeXt-Base pretrained for supervised classification on ImageNet-1K, implemented in torchvision using the IMAGENET1K_V1 weights^26^. Thus, the primary comparison held the ConvNeXt-Base backbone family constant. Additional vision-only comparisons were conducted using ResNet-101 with the IMAGENET1K_V2 weights^27^ and VGG-16 with the IMAGENET1K_V1 weights^28^.

All pretrained model parameters were held fixed throughout the analyses. For CLIP, we extracted the 512-dimensional image embedding features returned by the image encoder after its learned projection layer. For ConvNeXt-Base, we extracted the 1,024-dimensional representation after global average pooling and the final LayerNorm, immediately before the ImageNet classification layer. For ResNet-101, we extracted the 2,048-dimensional global-average-pooling output immediately before the classification layer. For VGG-16, we extracted the 4,096-dimensional representation after the second fully connected layer and ReLU, captured at the subsequent dropout layer (classifier.5), which was inactive in evaluation mode. Each embedding feature vector was subsequently normalized to unit Euclidean norm.

As a control analysis, we compared the vision–language and vision-only ConvNeXt-Base models using representations extracted from the same position within their visual backbones (see SI text 1 for results). For the vision–language model (CLIP), we extracted the globally pooled 1,024-dimensional representation immediately before the learned CLIP projection layer. For the vision-only model (ConvNeXt), we extracted the corresponding globally pooled 1,024-dimensional representation immediately before the ImageNet classification layer. Both representations were obtained from identically structured timm ConvNeXt-Base trunks. Image resizing, center cropping and interpolation were identical across models, whereas input normalization followed the original pretraining specification of each model.

Prediction analyses were conducted separately for each participant in the Online 1, Online 2 and fMRI datasets. Ridge regression models with an intercept were fitted to predict subjective value ratings from the image embedding features. The L2-regularization parameter was fixed at α = 1 for all models, participants and datasets, and no hyperparameter optimization was performed.

Generalization to held-out images was evaluated using five-fold cross-validation within each participant. Prediction accuracy in each fold was quantified as the Pearson correlation between predicted and observed ratings for the held-out images. The five fold-specific correlation coefficients were Fisher z-transformed and averaged, and the resulting mean was transformed back to the correlation scale to obtain one prediction-accuracy value for each participant and model.

Statistical analyses were performed on Fisher z-transformed participant-level prediction accuracies. Above-chance prediction by CLIP and ConvNeXt in the online datasets was assessed using two-tailed one-sample *t*-tests against zero. Bonferroni-adjusted *p* values were calculated across the eight planned tests comprising two models, two online datasets and two participant groups. Differences between CLIP and each vision-only CNN model were assessed using two-tailed paired-samples *t*-tests. For these analyses, Bonferroni-adjusted *p* values were calculated across the three planned CLIP-versus-CNN comparisons within each dataset and participant group.

#### Representational similarity analysis of model content

We used representational similarity analysis (RSA)^33^ to characterize the food-related information encoded by the vision-language (CLIP) and vision-only (ConvNeXt) models. For each model, we constructed model-based representational dissimilarity matrices (RDMs) across the food images using the embedding feature vectors described above. Dissimilarity between each pair of images was defined as one minus the Pearson correlation between their feature vectors.

Attribute-based RDMs were constructed from the RGB color values and food-pics extended annotations for the following properties: image statistics comprising contrast, normalized complexity and median spatial-frequency power; nutritional composition comprising protein, fat and carbohydrate content per 100 g; processing state (whole or processed); and taste category (sweet or tasty, following the database labels). Each continuous variable was standardized across images to have zero mean and unit variance before calculating Euclidean distances within each attribute set. For nutritional variables, means and standard deviations were calculated from non-missing observations, and missing values were retained without imputation. Processing-state and taste RDMs were calculated from binary category codes, yielding a distance of zero for images in the same category and one for images in different categories.

For each model–attribute combination, representational similarity was quantified as the Spearman rank correlation between the lower-triangular elements of the model and attribute RDMs, excluding the diagonal. Differences between models were evaluated using an image-level bootstrap with 10,000 resamples. In each iteration, 896 image indices were sampled with replacement, and the same indices were used to select rows and columns from both model RDMs and all attribute RDMs. Spearman correlations were recalculated from the lower triangles of the resampled matrices, and their difference was computed as CLIP minus ConvNeXt. The 2.5th and 97.5th percentiles of the bootstrap differences defined the 95% confidence interval. Two-sided bootstrap p-values were calculated as twice the smaller proportion of differences at or below zero and at or above zero, using a plus-one correction to both the count and the number of valid resamples and capping the resulting p-value at one.

We additionally examined representations of perceived healthiness using individual ratings from the Online 1 and Online 2 datasets. For each participant, a healthiness RDM was constructed using the absolute difference in ratings between each pair of images. The lower-triangular elements of this RDM were correlated with those of each model RDM using Spearman correlation. Model differences were assessed separately for each dataset and participant group using two-tailed paired t-tests on Fisher z-transformed participant-level RSA correlations.

#### Layer-wise representational analysis of food attributes

To examine how food-related information varied across the vision-language model, we extracted representations at six successive processing stages of the CLIP image encoder: the convolutional stem, the outputs of the four ConvNeXt stages, and the final image embedding. These representations corresponded to layers 1–6 in the figures. Intermediate activations were extracted from the pretrained model. For activations with spatial dimensions, we applied global average pooling across the height and width dimensions to obtain one feature vector per image. The final representation was the L2-normalized image embedding used in the prediction analyses.

For each layer, we constructed an RDM across the 896 food images using correlation distance, defined as one minus the Pearson correlation between feature vectors. We then calculated Spearman correlations between the lower-triangular elements of each layer RDM and the attribute RDMs for color, image statistics, nutritional composition, processing state and taste category. Each layer was analyzed independently, yielding one representational-similarity estimate per layer and attribute set.

We also examined the correspondence of layer representations with perceived healthiness and subjective value using individual ratings. For each participant and rating dimension, we constructed an RDM from the absolute differences between ratings for pairs of images. We correlated its lower-triangular elements with those of each layer RDM using Spearman correlation. Participant-level correlations were summarized as the mean and standard error of the untransformed coefficients, separately for each dataset and participant group.

### fMRI data acquisition

Functional MRI images were collected using a 3 T Siemens MAGNETOM Prisma scanner at the HIAS Brain Research Center (Hitotsubashi University, Tokyo, Japan) with a 32-channel radio frequency coil. The BOLD signal was measured using a T2*-weighted gradient-echo echo-planar imaging (EPI) sequence with the following parameters: TR = 800 ms, TE = 34.4 ms, flip angle = 52°, voxel size = 2.4 × 2.4 × 2.4 mm, matrix size = 86 × 86, 60 slices aligned to the AC–PC plane, and multiband acceleration factor = 6. For susceptibility distortion correction of the fMRI images, we also acquired a pair of spin-echo EPI “fieldmap” images with opposite phase-encoding directions (anterior-to-posterior [AP] and posterior-to-anterior [PA]) using the same geometry (FOV, matrix size, slice orientation and thickness) as the functional runs. High-resolution (0.9 mm^3^) anatomical images were acquired using a standard MPRAGE pulse sequence (TR = 2,300 ms, TE = 2.32 ms, FA = 8°).

### fMRI data analysis

#### Preprocessing

fMRI data preprocessing was performed using fMRIPrep 25.2.3^52^, which is based on Nipype 1.10.0. The fMRIPrep pipeline included intensity non-uniformity correction, skull stripping, head motion correction, susceptibility distortion correction using fieldmaps, co-registration of functional and anatomical images, and spatial normalization to the MNI152NLin2009cAsym template.

#### Univariate analysis

We performed univariate fMRI analysis using general linear models implemented in SPM25. Spatially normalized functional images were smoothed with a Gaussian kernel of 4 mm full width at half maximum (FWHM). A separate design matrix was specified for each run within each participant. In the model, the valuation phase was modeled as a boxcar regressor from image onset to the onset of the rating phase, with a duration of 4 s, and included a first-order parametric modulator encoding the subjective value rating on each trial. Additional regressors modeled the rating phase, from rating-screen onset to the button response; the button response as a zero-duration event; and the feedback phase as a 0.5-s boxcar. Missed trials were modeled separately. Those regressors were convolved with the canonical hemodynamic response function. Six head-motion parameters and indicator regressors for volumes with framewise displacement exceeding 0.9 mm were included as nuisance regressors. Low-frequency drifts were removed using a high-pass filter with a 128-s cutoff, and temporal autocorrelation was modeled using a first-order autoregressive process.

For each participant, we constructed a contrast for the subjective-value parametric modulator across runs. Participant-level contrast images were entered into a group-level analysis. Whole-brain results were assessed at a voxel-level family-wise error (FWE)-corrected threshold of *p* < 0.05.

We also examined subjective-value effects in independently defined regions of interest (ROIs) implicated in food valuation^12^. Spherical ROIs with an 8-mm radius were centered at MNI coordinates [−2, 40, −6] for the ventromedial prefrontal cortex, [−10, 8, −6] for the left ventral striatum, and [10, 12, −10] for the right ventral striatum. These coordinates were obtained from a previous meta-analysis^12^.

#### Multivoxel pattern analysis

We used representational similarity analysis (RSA), implemented in CoSMoMVPA^53^, to examine the correspondence between model embedding features and multivoxel fMRI activity patterns during food valuation. Trial-specific activity patterns were estimated from unsmoothed functional images using a GLM approach. Each image presentation was modeled with its own valuation-phase regressor, allowing a separate beta image to be estimated for each trial. Additional regressors modeled the rating phase, button responses and feedback. The canonical hemodynamic response function, motion nuisance regressors, and high-pass filtering followed the settings described above. Trial-specific beta images for the valuation-phase were then fed into RSA.

Whole-brain RSA used spherical searchlights with a radius of three voxels. Within each searchlight, a neural RDM was calculated using correlation distance between trial-specific activity patterns. Model RDMs were constructed using correlation distance between the corresponding image features from the vision-language (CLIP) and vision-only (ConvNeXt) model. A subjective-value RDM was constructed separately for each participant using absolute differences between subjective value ratings. All neural RSA analyses used only image pairs presented in different runs to reduce the influence of shared within-run noise.

We first calculated Spearman correlations between the neural RDM and each model RDM separately. Subjective-value RSA was conducted using the same cross-run procedure. Correlation coefficients were Fisher z-transformed before group-level analyses. To directly compare the neural correspondence of the two models, we also conducted multiple-regression RSA^35,36^, in which we regressed fMRI-based RDMs on RDMs derived from the vision-language and vision-only models.

To examine layer-dependent neural correspondence, we repeated the searchlight RSA independently for the six CLIP layers described above. We further performed multiple-regression RSA^35,36^ to estimate the unique contributions of selected layers and subjective value. Because adjacent layers showed strongly correlated representational structures, this analysis included the RDMs for layers 1, 4 and 6, together with the participant-specific subjective-value RDM, as simultaneous predictors of the neural RDM. The resulting regression coefficients quantified each predictor’s contribution conditional on the other predictors and were entered into group-level analyses.

Participant-level searchlight maps were smoothed with a 4-mm FWHM Gaussian kernel and entered into group-level one-sample t-tests in SPM25. Positive effects were assessed at *p* < 0.05 with voxel-level FWE correction.

We also performed RSA within independently defined anatomical ROIs. Visual ROIs were derived from the Wang retinotopic atlas^54^, and downstream temporal, parietal and prefrontal ROIs were defined using the Brainnetome atlas^55^. Dorsal and ventral subdivisions of V1–V3 were merged within each hemisphere. VO1 and VO2, PHC1 and PHC2, LO1 and LO2, and IPS0 and IPS1 were likewise combined into VO, PHC, LO and IPS ROIs, respectively. RSA was calculated from the multivoxel patterns within each ROI using the same RDM definitions, cross-run pair selection and Spearman or partial Spearman correlations as in the searchlight analyses. Correlations were Fisher z-transformed, and values from homologous left and right ROIs were averaged within each participant.

## Data availability

The data are available at https://osf.io/krgh3/files/osfstorage (Online 1), https://osf.io/puqsh/files/osfstorage (Online 2) and https://openneuro.org/datasets/ds007267/versions/1.1.1 (fMRI).

## Author contributions

A.T., Y.Y. and S.S. designed the experiments. A.T., M.S., Y.A., Y.Y. and S.S. collected the data. Y.A, A.T., H.K. and S.S. analyzed the data. S.S. supervised the research. S.S. drafted the manuscript. All authors reviewed and approved the final manuscript.

## Supplementary materials

### SI text 1: Prediction accuracy of the vision-language and vision-only models using matched backbone representations

The controlled analysis replicated the pattern observed in the primary analysis (Fig. 1b). When both models were represented by 1,024-dimensional features extracted from the corresponding pre-head position of the ConvNeXt-Base backbone, prediction accuracy remained higher for the vision-language model than for the vision-only model (Online 1: lean, *t*(105) = 28.119, *p* < 0.001; obese, *t*(87) = 22.401, *p* < 0.001; Online 2: lean, *t*(106) = 20.667, *p* < 0.001; obese, *t*(92) = 18.588, *p* < 0.001). Thus, the advantage of the vision-language model could not be attributed solely to differences in feature dimensionality or to comparing features extracted from different positions relative to the task-specific output heads.

**Table S1.** Clusters showing significant correspondence between model embedding features (i.e., activity pattern of the output layer) and multivoxel fMRI activity patterns.

| **Model** | **Region** | **x** | **y** | **z** | **t-statistic** | ***p*<sub>FWE</sub>** | **Voxels** |
|---|---|---|---|---|---|---|---|
| Vision-language | Right fusiform gyrus | 36 | -60 | -11 | 11.65 | <0.001 | 10,791 |
|  | Right fusiform gyrus | 38 | -22 | -25 | 7.96 | <0.001 | 36 |
|  | Left middle frontal gyrus | -43 | 33 | 25 | 7.57 | <0.001 | 57 |
|  | Right precentral gyrus | 53 | 2 | 28 | 6.38 | 0.009 | 19 |
|  | Right postcentral gyrus | 55 | -24 | 42 | 5.98 | 0.023 | 18 |
|  | Left angular gyrus | -51 | -67 | 30 | 5.85 | 0.030 | 16 |
| Vision-only | Right inferior occipital gyrus | 33 | -84 | -8 | 13.26 | <0.001 | 11,484 |
|  | Left inferior parietal lobule | -50 | -41 | 52 | 7.25 | 0.002 | 57 |
|  | Right inferior parietal lobule | 45 | -39 | 49 | 6.68 | 0.006 | 82 |
| Vision-language vs. Vision-only | Left fusiform gyrus | -34 | -60 | -11 | 7.33 | <0.001 | 275 |
|  | Right fusiform gyrus | 38 | -70 | -8 | 6.85 | 0.002 | 99 |
|  | Right middle occipital gyrus | 43 | -84 | 6 | 6.62 | 0.003 | 141 |
|  | Left middle occipital gyrus | -31 | -96 | 16 | 6.44 | 0.005 | 35 |

Whole-brain searchlight RSA results are reported at *p* < 0.05, voxel-level family-wise error corrected, with an additional extent threshold of k = 10 voxels.

**Table S2.** Clusters showing significant correspondence between each layer in the vision-language model and multivoxel fMRI activity patterns.

| **Layer** | **Region** | **x** | **y** | **z** | **t-statistic** | ***p*<sub>FWE</sub>** | **Voxels** |
|---|---|---|---|---|---|---|---|
| 1 | Left lingual gyrus | -15 | -86 | -8 | 12.22 | <0.001 | 5,224 |
| 2 | Left lingual gyrus | -17 | -89 | -11 | 11.92 | <0.001 | 5,902 |
|  | Right superior occipital gyrus | 24 | -63 | 47 | 6.15 | 0.011 | 13 |
| 3 | Left middle occipital gyrus | -17 | -94 | 4 | 10.90 | <0.001 | 4,950 |
|  | Right superior occipital gyrus | 24 | -63 | 47 | 7.17 | 0.001 | 35 |
| 4 | Left lingual gyrus | -17 | -82 | -11 | 9.69 | <0.001 | 2,595 |
| 5 | Left inferior occipital gyrus | -41 | -77 | -6 | 13.28 | <0.001 | 11,212 |
|  | Right precentral gyrus | 33 | 0 | 44 | 7.09 | 0.002 | 26 |
|  | Right middle frontal gyrus | 41 | 36 | 20 | 7.04 | 0.002 | 52 |
|  | Left precentral gyrus | -53 | 7 | 32 | 6.37 | 0.012 | 25 |
|  | Right precentral gyrus | 57 | 7 | 32 | 6.12 | 0.021 | 13 |
| 6 | Right fusiform gyrus | 36 | -60 | -11 | 11.65 | <0.001 | 10,791 |
|  | Right fusiform gyrus | 38 | -22 | -25 | 7.96 | <0.001 | 36 |
|  | Left middle frontal gyrus | -43 | 33 | 25 | 7.57 | <0.001 | 57 |
|  | Right precentral gyrus | 53 | 2 | 28 | 6.38 | 0.009 | 19 |
|  | Right postcentral gyrus | 55 | -24 | 42 | 5.98 | 0.023 | 18 |
|  | Left angular gyrus | -51 | -67 | 30 | 5.85 | 0.030 | 16 |

Whole-brain searchlight RSA results are reported at *p* < 0.05, voxel-level family-wise error corrected, with an additional extent threshold of k = 10 voxels.

**Table S3.**
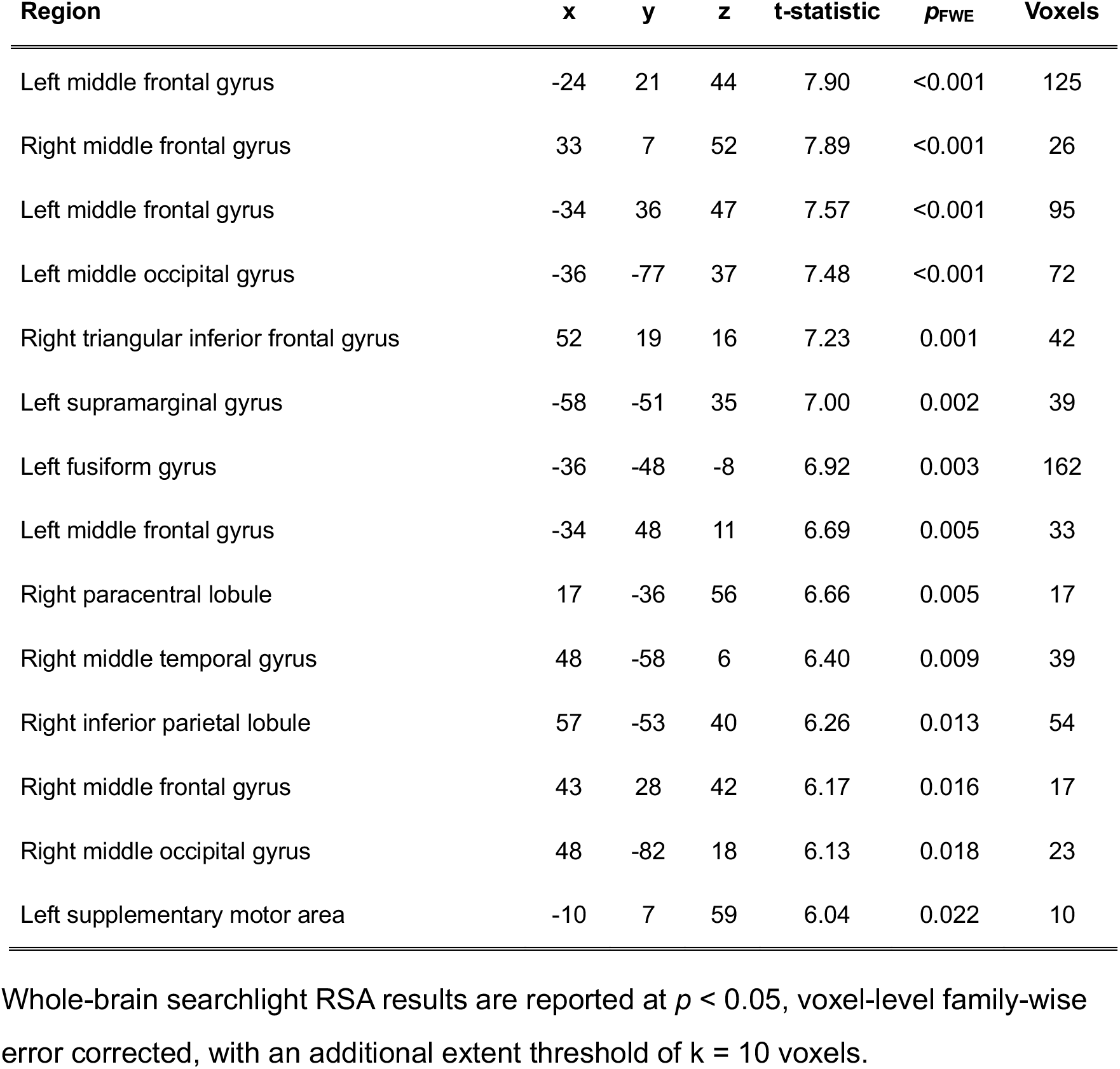
Clusters showing significant correspondence between vision-language model-predicted subjective value and multivoxel fMRI activity patterns.

| **Region** | **x** | **y** | **z** | **t-statistic** | ***p*<sub>FWE</sub>** | **Voxels** |
|---|---|---|---|---|---|---|
| Left middle frontal gyrus | -24 | 21 | 44 | 7.90 | <0.001 | 125 |
| Right middle frontal gyrus | 33 | 7 | 52 | 7.89 | <0.001 | 26 |
| Left middle frontal gyrus | -34 | 36 | 47 | 7.57 | <0.001 | 95 |
| Left middle occipital gyrus | -36 | -77 | 37 | 7.48 | <0.001 | 72 |
| Right triangular inferior frontal gyrus | 52 | 19 | 16 | 7.23 | 0.001 | 42 |
| Left supramarginal gyrus | -58 | -51 | 35 | 7.00 | 0.002 | 39 |
| Left fusiform gyrus | -36 | -48 | -8 | 6.92 | 0.003 | 162 |
| Left middle frontal gyrus | -34 | 48 | 11 | 6.69 | 0.005 | 33 |
| Right paracentral lobule | 17 | -36 | 56 | 6.66 | 0.005 | 17 |
| Right middle temporal gyrus | 48 | -58 | 6 | 6.40 | 0.009 | 39 |
| Right inferior parietal lobule | 57 | -53 | 40 | 6.26 | 0.013 | 54 |
| Right middle frontal gyrus | 43 | 28 | 42 | 6.17 | 0.016 | 17 |
| Right middle occipital gyrus | 48 | -82 | 18 | 6.13 | 0.018 | 23 |
| Left supplementary motor area | -10 | 7 | 59 | 6.04 | 0.022 | 10 |

Whole-brain searchlight RSA results are reported at *p* < 0.05, voxel-level family-wise error corrected, with an additional extent threshold of k = 10 voxels.

**Table S4.**
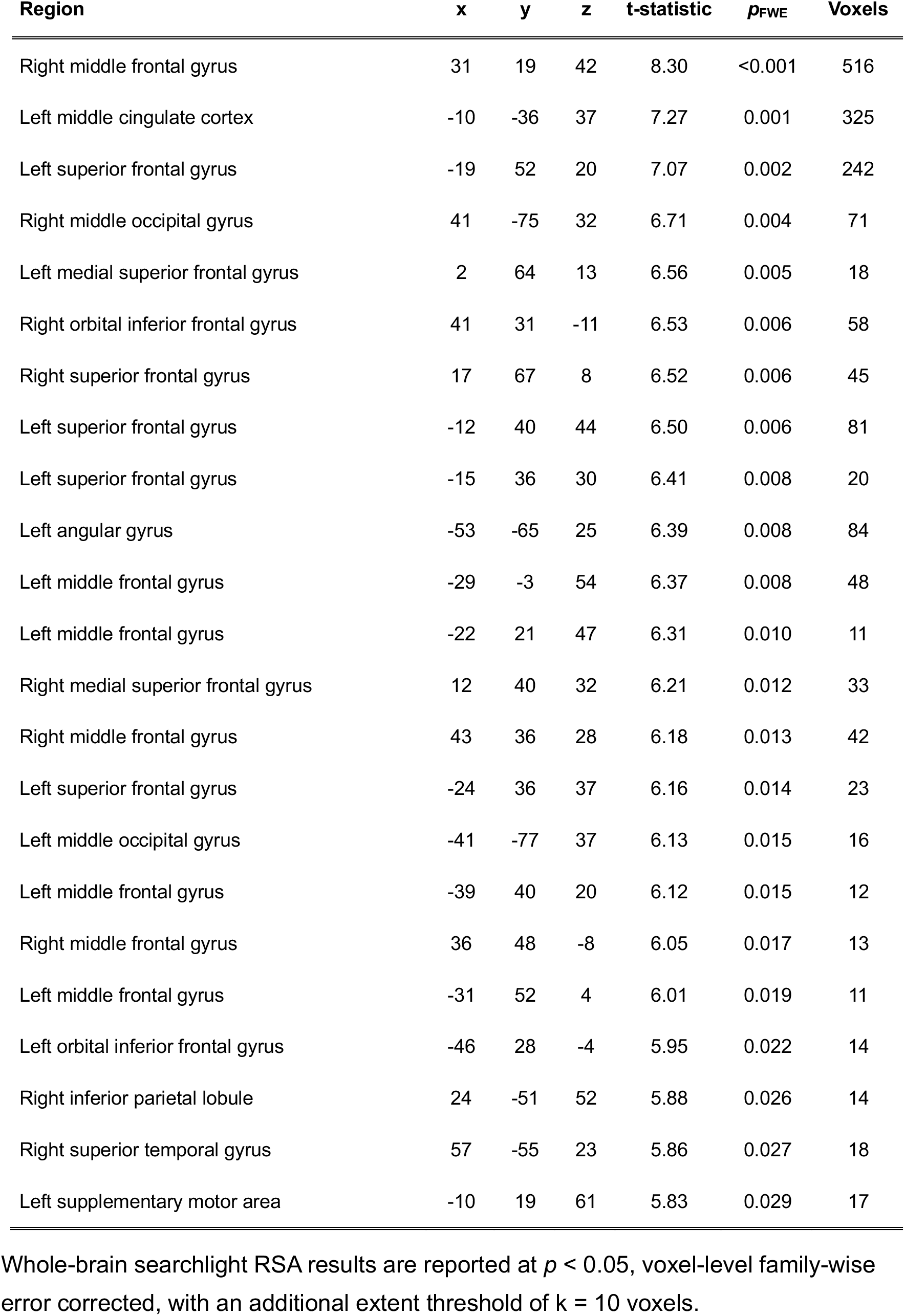
Clusters showing significant correspondence between subjective value rating and multivoxel fMRI activity patterns.

**Figure S1:**
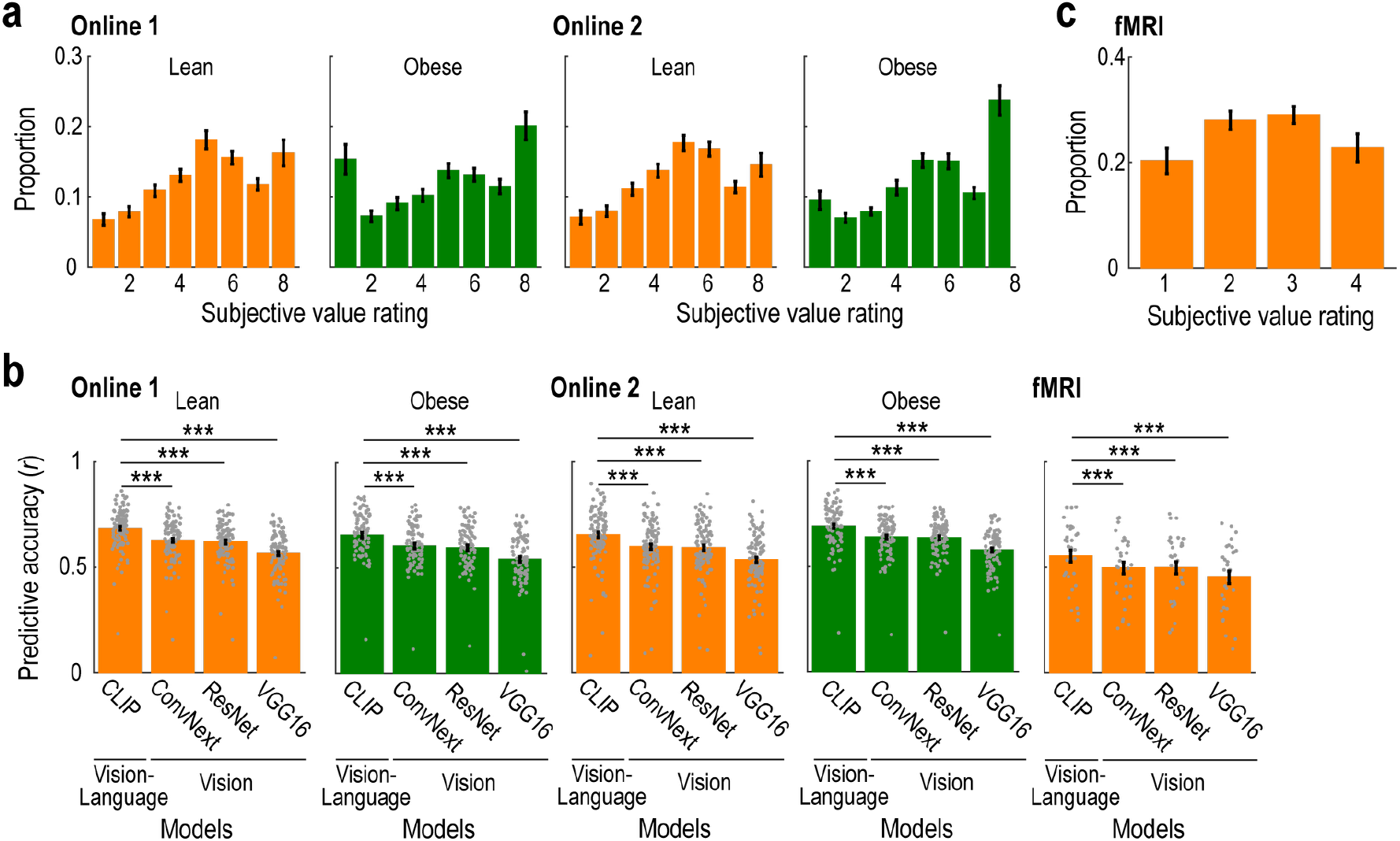
Rating distributions and supplementary analyses of prediction performance. (a) Rating distributions in the Online 1 and Online 2 experiments (mean ± SEM across participants). (b) Prediction accuracy, quantified as the cross-validated correlation between predicted and observed ratings (mean ± SEM across participants). Each grey dot represents an individual participant. \*\*\**p* < 0.001, *t*-test (two-tailed). (c) Rating distributions in the fMRI experiment. The format is the same as in (a).

**Figure S2:**
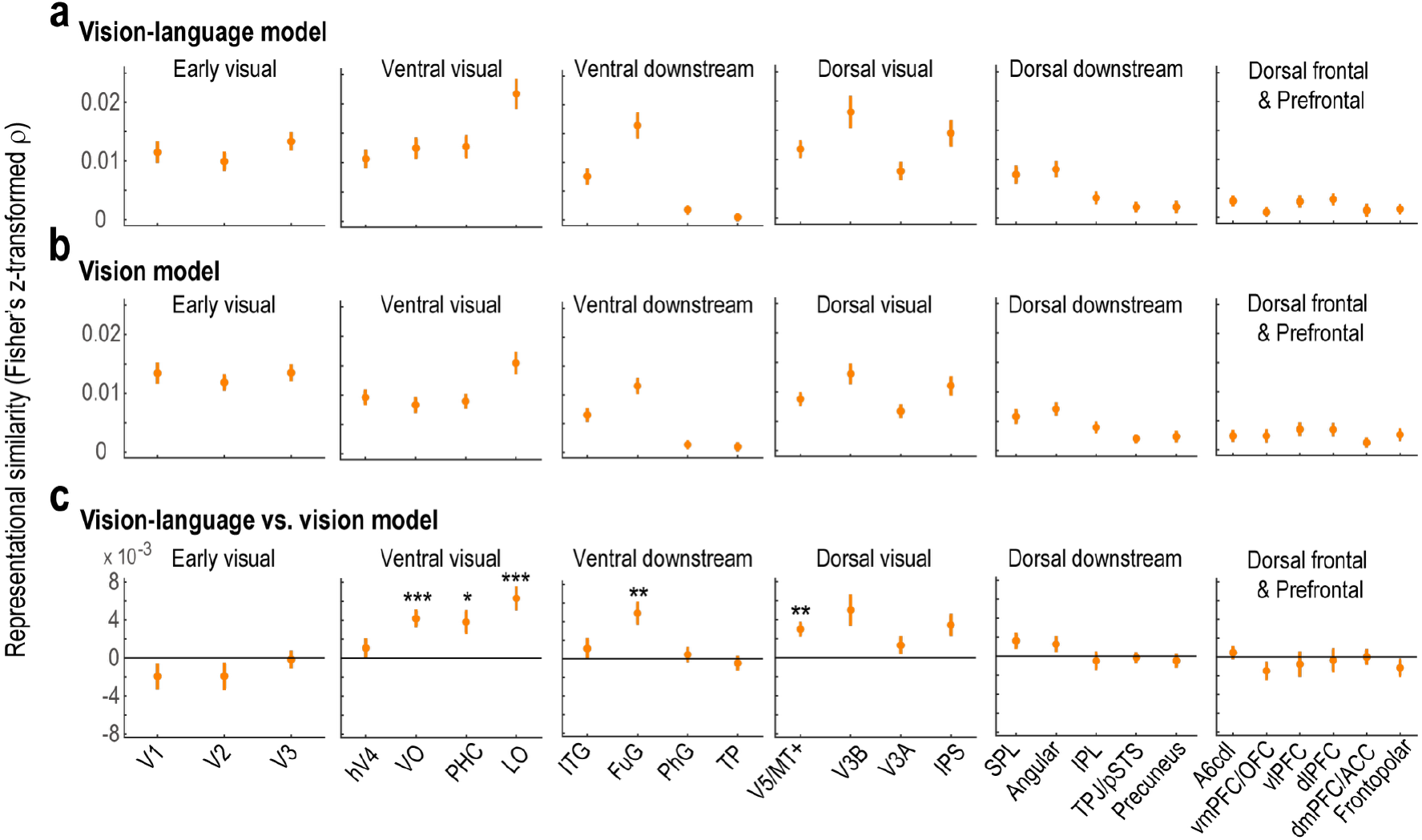
Anatomically defined ROI analysis of the neural representations of model embedding features. (a) Representational similarity between embedding features from the vision-language models and multivoxel fMRI activity patterns during the valuation phase for each anatomically defined region of interest (ROI) (mean ± SEM across participants). V1, Primary visual cortex / primary visual area; V2, Secondary visual cortex / visual area 2; V3, Visual area 3; hV4, Human visual area 4; VO, Ventral occipital areas; PHC, Parahippocampal cortex; LO, Lateral occipital areas; ITG, Inferior temporal gyrus; FuG, Fusiform gyrus; PhG, Parahippocampal gyrus; Temporal pole, Temporal pole; V5/MT+, Visual area 5 / middle temporal visual complex; V3B, Visual area V3B; V3A, Visual area V3A; IPS, Intraparietal sulcus; SPL, Superior parietal lobule; Angular, Angular gyrus; IPL, Inferior parietal lobule; TPJ/pSTS, Temporoparietal junction / posterior superior temporal sulcus; A6cdl, Caudodorsal lateral area 6 (dorsal premotor cortex); vmPFC/OFC, Ventromedial prefrontal cortex / orbitofrontal cortex; vlPFC, Ventrolateral prefrontal cortex; dlPFC, Dorsolateral prefrontal cortex; dmPFC/ACC, Dorsomedial prefrontal cortex / anterior cingulate cortex; Frontopolar, Frontopolar cortex. (b) Representational similarity between embedding features from the vision-only models and multivoxel fMRI activity patterns during the valuation phase for each anatomically defined ROI (mean ± SEM across participants). The format is the same as in (a). (c) Difference between the vision-language and vision-only models in the representational similarity with multivoxel fMRI activity patterns for each anatomically ROI (mean ± SEM across participants). The format is the same as in (a). \**p* < 0.05, \*\**p* < 0.01, and \*\*\**p* < 0.001 Bonferroni-corrected across 26 ROIs, *t*-test (two-tailed).

**Figure S3:**
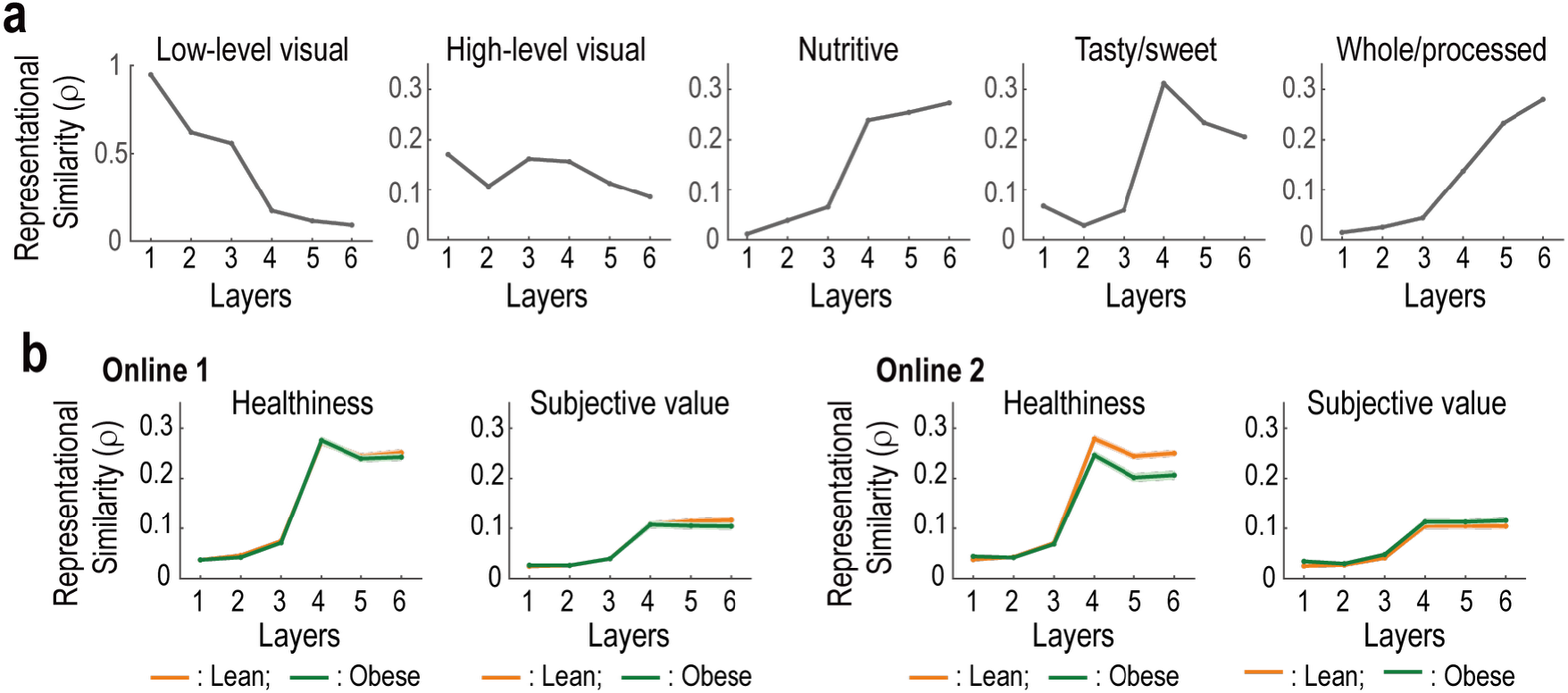
Supplementary analysis of the layer-wise representations in the vision-language model. (a) Representation of visual, nutritional, taste and processing-state attributes across layers. For each layer, we plotted representational similarity, quantified as Spearman’s rank correlation (ρ), between the model embedding features and each food attribute. Note that the embedding features were extracted from each layer after normalization. (b) Representation of perceived healthiness across layers. For each layer, we plotted representational similarity between the model embedding features and perceived healthiness ratings as well as subjective value (mean± SEM across participants). *Left*, Online 1. *Right*, Online 2.

**Figure S4:**
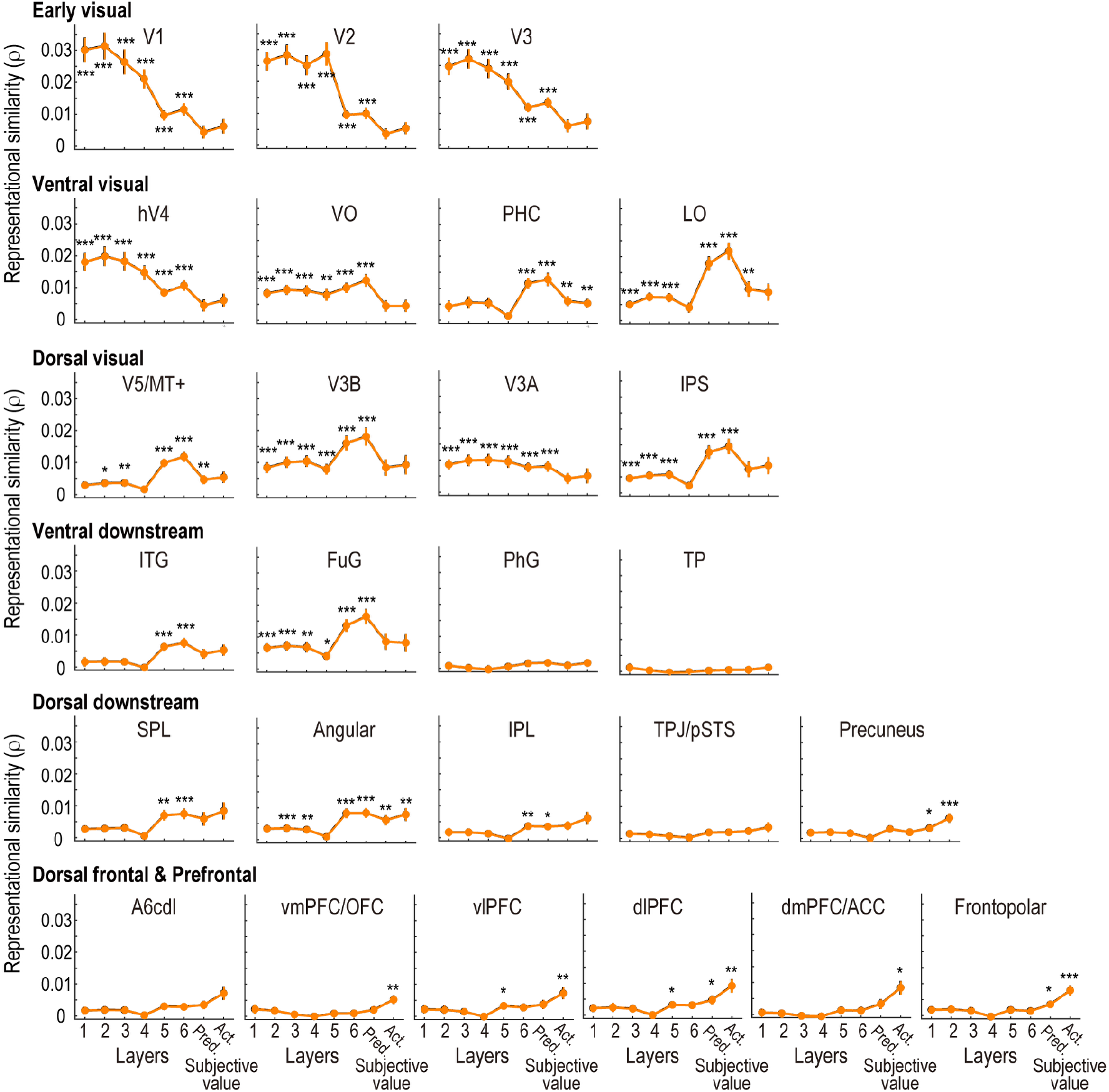
Supplementary anatomically-defined ROI analysis of the layer-wise neural representations in the vision-language model. Representational similarity between embedding features from each layer in the vision-language model and multivoxel fMRI activity patterns during the valuation phase is plotted for each anatomically defined region of interest (ROI). Points and error bars indicate the mean ± SEM across participants. V1, Primary visual cortex / primary visual area; V2, Secondary visual cortex / visual area 2; V3, Visual area 3; hV4, Human visual area 4; VO, Ventral occipital areas; PHC, Parahippocampal cortex; LO, Lateral occipital areas; ITG, Inferior temporal gyrus; FuG, Fusiform gyrus; PhG, Parahippocampal gyrus; Temporal pole, Temporal pole; V5/MT+, Visual area 5 / middle temporal visual complex; V3B, Visual area V3B; V3A, Visual area V3A; IPS, Intraparietal sulcus; SPL, Superior parietal lobule; Angular, Angular gyrus; IPL, Inferior parietal lobule; TPJ/pSTS, Temporoparietal junction / posterior superior temporal sulcus; A6cdl, Caudodorsal lateral area 6 (dorsal premotor cortex); vmPFC/OFC, Ventromedial prefrontal cortex / orbitofrontal cortex; vlPFC, Ventrolateral prefrontal cortex; dlPFC, Dorsolateral prefrontal cortex; dmPFC/ACC, Dorsomedial prefrontal cortex / anterior cingulate cortex; Frontopolar, Frontopolar cortex. Pred, predicted subjective value rating. Act., actual subjective value rating. \**p* < 0.05, \*\**p* < 0.01, and \*\*\**p* < 0.001 Bonferroni-corrected across 26 ROIs × [6 layers + predicted + actual subjective value ratings], *t*-test (two-tailed).

**Figure S5:**
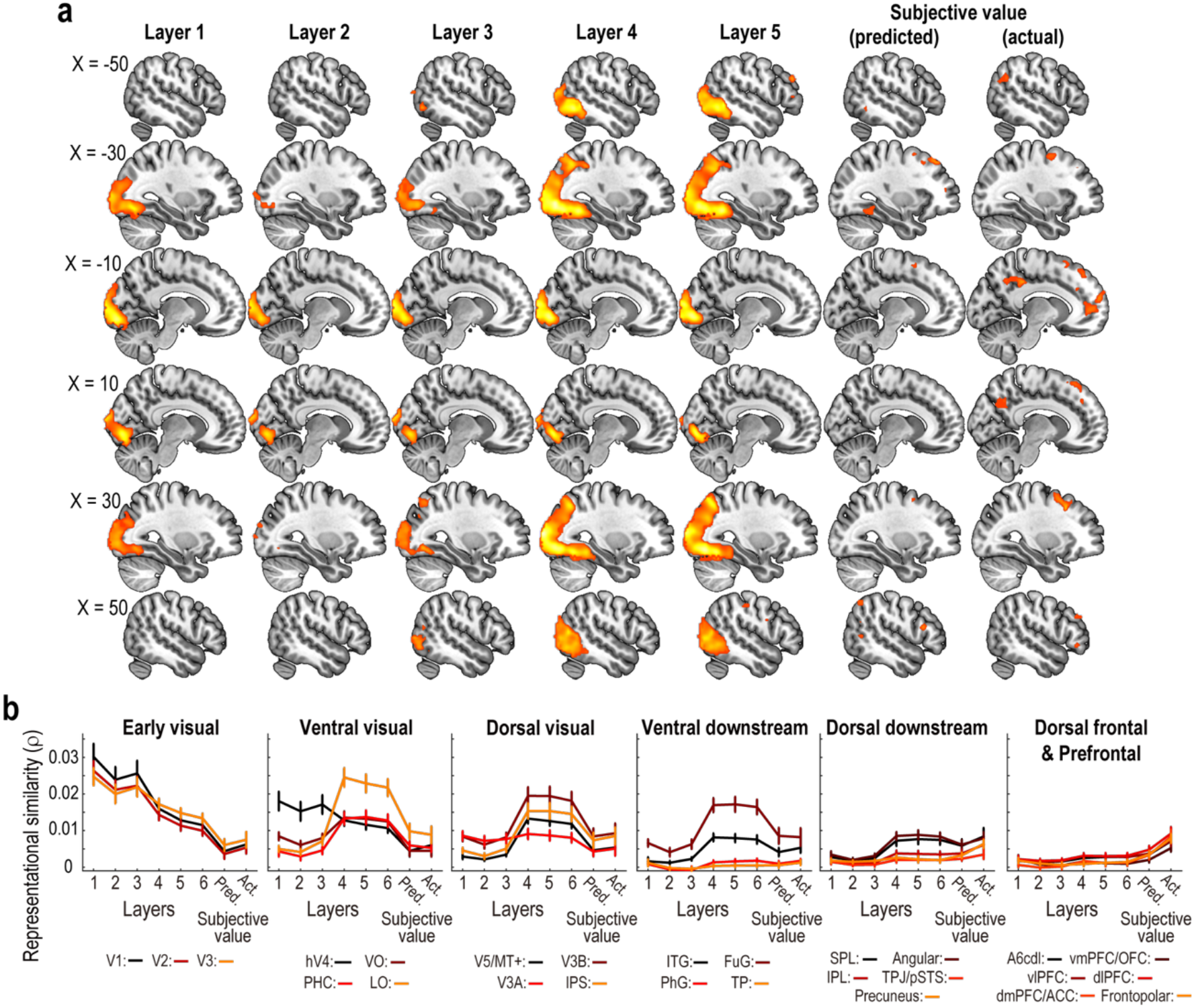
Supplementary analysis of the layer-wise neural representations in the vision-language model. (a) Whole-brain searchlight representational similarity analysis (RSA) of each layer in the vision-language model. The format is the same as Fig. 6a. Note that the embedding features were extracted from each layer after normalization. (b) Anatomically defined ROI RSA of each layer in the vision-language model and the subjective values. The format is the same as Fig. 6b.

**Figure S6:**
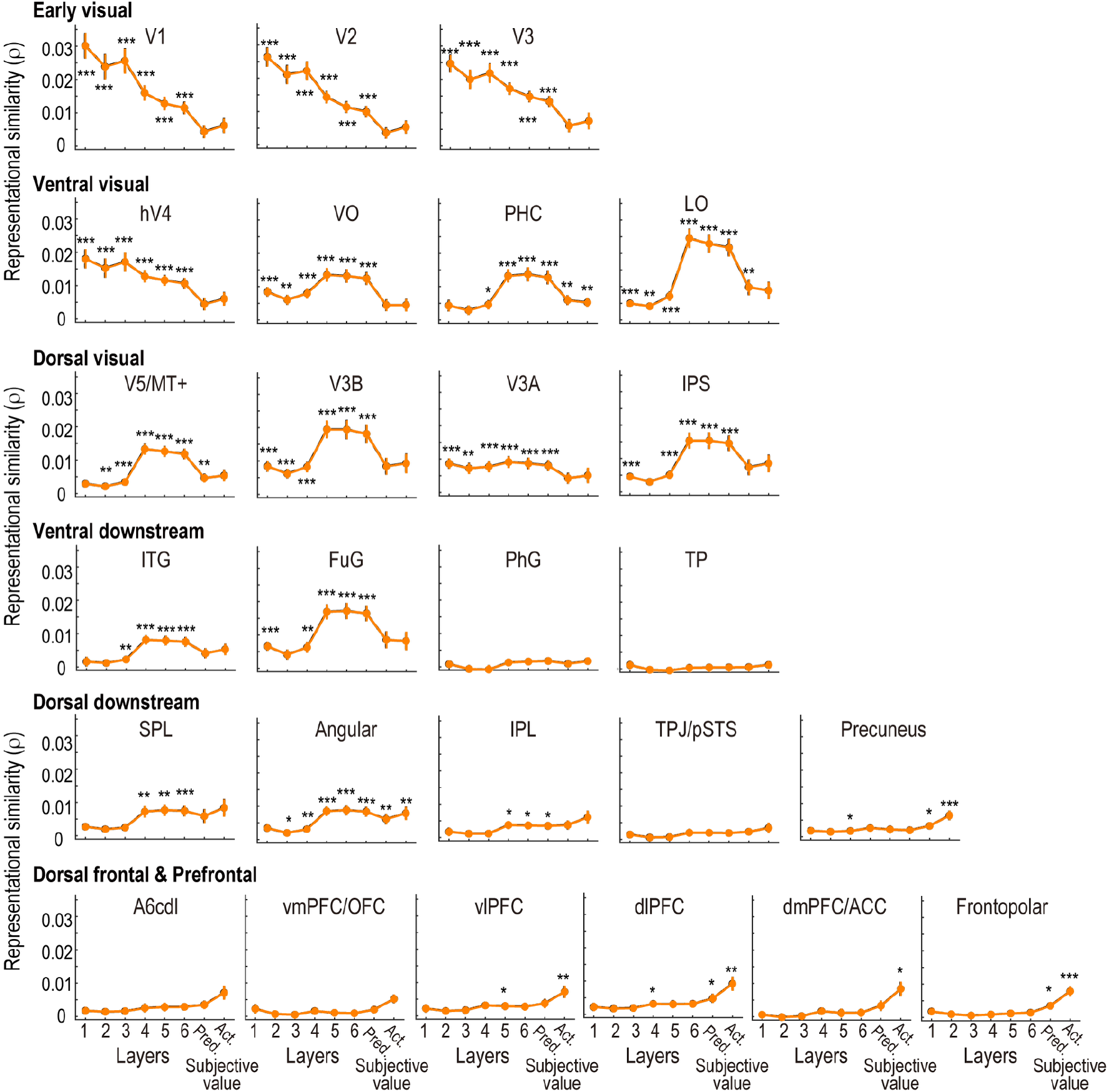
Supplementary anatomically-defined ROI analysis of the layer-wise neural representations in the vision-language model. Representational similarity between embedding features from each layer in the vision-language model and multivoxel fMRI activity patterns during the valuation phase is plotted for each anatomically defined region of interest (ROI). Note that the embedding features were extracted from each layer after normalization. The format is the same as Fig. S4.

**Figure S7:**
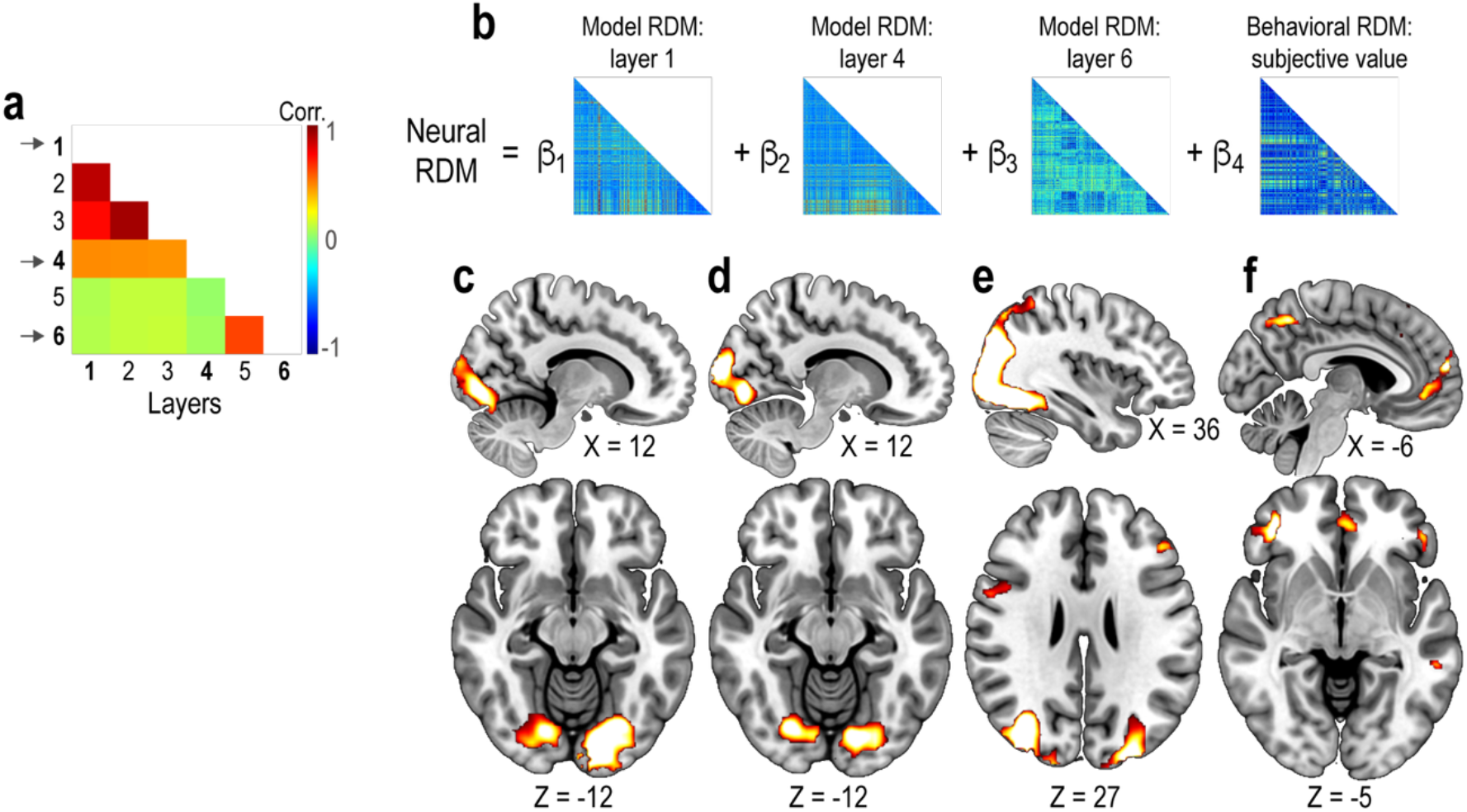
Supplementary results on the layer-wise neural representations in the vision-language model. (a) Cross-correlation matrix of the embedding features across layers of the vision-language model. Arrows indicate the layers used in the multiple-regression representational similarity analysis (RSA). (b) Schematic illustration of the multiple-regression RSA. (c) Whole-brain searchlight RSA of layer 1 in the vision-language model. Statistical parametric maps show representational similarity between model embedding features and multivoxel fMRI activity patterns during the valuation phase. The maps are thresholded at *p* < 0.05, voxel-level FWE corrected. (d) Whole-brain searchlight RSA of layer 4. The format is the same as (c). (e) Whole-brain searchlight RSA of layer 6. The format is the same as (c). (f) Whole-brain searchlight RSA of subjective value. The format is the same as (c).

